# CK1a-Mediated Two-Step Subunit Remodeling of the Circadian FRQ-FRH Complex

**DOI:** 10.1101/2025.05.13.653691

**Authors:** Carolin Schunke, Bianca Ruppert, Sabine Schultz, Axel C.R. Diernfellner, Michael Brunner

## Abstract

The circadian clock of *Neurospora* operates through a negative feedback loop in which FRQ, along with FRH and CK1a, inhibits its transcriptional activator, WCC, via phosphorylation. CK1a, anchored to FRQ, hyperphosphorylates FRQ at its IDRs in a slow, temperature-independent manner, forming a module suited for molecular timekeeping. However, the molecular processes triggered by FRQ’s hyperphosphorylation have remained unclear. We show that FRH, the folded binding partner of disordered FRQ, decodes FRQ’s time-dependent phosphorylation state by triggering a two-step remodeling of the FRQ-FRH complex: initially, two FRH molecules bind a FRQ dimer, keeping it inactive by blocking its interaction with WCC. Gradual phosphorylation induces with a delay the dissociation of one FRH, exposing a binding site for WCC and activating the complex. Following a time delay, attributable to the slow and stochastic nature of phosphorylation, the release of the second FRH promotes nuclear export and subsequent degradation of FRQ. This stepwise remodeling ensures precise activation and inactivation of FRQ and positions FRH as a hub for decoding temporal phosphorylation information.

## Introduction

Circadian clocks are fascinating molecular devices capable of accurately measuring time and orchestrating the physiology and behavior of organisms in sync with the 24-hour environmental day-night cycle. The circadian clock of the filamentous fungus *Neurospora crassa* is based on a delayed transcriptional-translational negative feedback loop: The heterodimeric circadian transcription factor White Collar Complex (WCC), composed of a White Collar-1 (WC-1) and WC-2 subunit, activates transcription of the circadian clock gene *frequency* (*frq*) ^1,2^. The FRQ protein, 85% of which consists of intrinsically disordered regions (IDRs)^2^, dimerizes via a coiled-coil in its N-terminal region ^3^. FRQ-Interacting RNA-helicase, FRH, binds to the C-terminal portion of FRQ and protects FRQ from premature degradation ^4^. FRH is an orthologue of the conserved MTR-4 helicase ^4^, which has essential functions in various aspects of RNA metabolism ^5^. Helicase activity of FRH is not required for its moonlighting function in the circadian clock ^6^. Casein kinase 1a (CK1a) binds to the central part of FRQ via the spatially separated FRQ-CK1-interaction Domains 1 and 2, FCD1 and FCD2 ^7,8^. The FRQ-FRH-CK1a complex (FFC) interacts weakly and dynamically with WCC and facilitates its inactivation through phosphorylation by CK1a ^9^. In the course of a circadian period FRQ is progressively hyperphosphorylated leading to its inactivation and degradation ^10–12^. More than 100 phosphorylation sites in FRQ have been identified ^13,14^, the vast majority located in its IDRs ^13,15,16^. Hyperphosphorylation is associated with circadian timekeeping and triggers a poorly characterized conformational change in the FFC ^8,17^. FCD-bound CK1a is sufficient to phosphorylate FRQ on a circadian timescale and in a temperature-insensitive manner, and thus the FFC constitutes a molecular module suited to measure time ^16,18^. Complete compensation of the clock at high temperature requires additionally phosphorylation of FRQ by casein kinase 2 ^19^.

Despite extensive knowledge of the components of the core clock and their interactions, it is still unclear how time is measured at the molecular level, which is the essential function of a circadian clock. This also applies to the circadian clocks of metazoans, vertebrates and invertebrates.

For example, it is not yet known how FRH protects FRQ from degradation, or whether this is its sole function. Furthermore, it is not known when and how the FFC interacts with WCC. Most significantly, we do not understand what kind of conformational change could be triggered by the partially redundant multisite phosphorylation of IDRs in FRQ and how this allows the FFC to perform its functions at the right time.

Answering these questions in *Neurospora* is a technical challenge and in many cases not possible. For example, CK1a and FRH have essential functions outside the clock and cannot be deleted or inactivated. FRQ and WC-1 without their binding partners, FRH and WC-2, respectively, are not expressed at a sufficient level to pursue complex biochemical analyses. Furthermore, *Neurospora* grows as a syncytium and forms dense hyphal networks with many nuclei that are rapidly transported along the hyphae ^20^, making live-cell microscopy challenging. In addition, the WCC is a light receptor that resets the circadian clock in response to blue light stimuli. Therefore, analyzing the subcellular distribution and dynamics of clock proteins by imaging in living cells is technically hardly feasible. These systemic and technical difficulties have greatly slowed down new insights into the functioning of the *Neurospora* circadian clock in recent years.

To overcome these challenges, we opted to heterologously express fluorescently tagged *Neurospora* clock proteins in U2OStx cells, enabling us to analyze their interactions and subcellular dynamics using a live cell imaging system. This straightforward approach enabled us to explore processes that are challenging or impossible to study in a cell-free *in vitro* system or in *Neurospora*. Based on these observations, we were then able to design specific experiments for validation in *Neurospora* that we might not have originally considered.

The approach yielded fundamentally new and unexpected insights into the timekeeping mechanisms of the core clock. Our data reveal that the phosphorylation state of FRQ’s intrinsically disordered regions (IDRs) is decoded through its interaction with its folded partner, FRH. FRH blocks the nuclear export signal (NES) of FRQ and the binding site for WCC, initially allowing accumulation of inactive inhibitor complexes that are activated with a delay by slow phosphorylation of FRQ. Phosphorylation eventually triggers the sequential stepwise release of FRH molecules. FFC subunit remodeling regulates FRQ’s delayed interaction with WCC, as well as its nuclear export and degradation.

## Results

### FRQ contains three nuclear localization signals

To analyze the subcellular localization of FRQ we constructed plasmids expressing full-length and truncated FRQ versions with either an N-terminal mKate2 (mK2) or a C-terminal mNeonGreen (mNG) moiety under control of a tetracycline-inducible CMV promoter (Fig. 1A, S1A). The proteins were heterologously expressed in U2OStx cells via transient transfection, and the fluorescently tagged proteins were analyzed using an Incucyte live-cell imaging system. Overexpressed mK2-FRQ and FRQ-mNG accumulated in nuclear foci (Fig. 1B left panels, S1B). FRQ^6B2^ denotes a DHF to AAA substitution of amino acid residues 774-776 in FRQ, which is part of a larger binding site and abolishes binding of FRH^21,22^, while FRQ^9^ is a protein truncated after aa 662 ^23^, entirely lacking the FRH binding site (Fig. 1A). mK2-FRQ^6B2^ formed nuclear foci like full-length mK2-FRQ and mK2-FRQ^9^ formed nuclear foci, which were slightly smaller (Fig. 1B). The data suggest that FRQ has the potential to self-interact and thus forms thermodynamically stable foci when overexpressed.

**Fig. 1.**
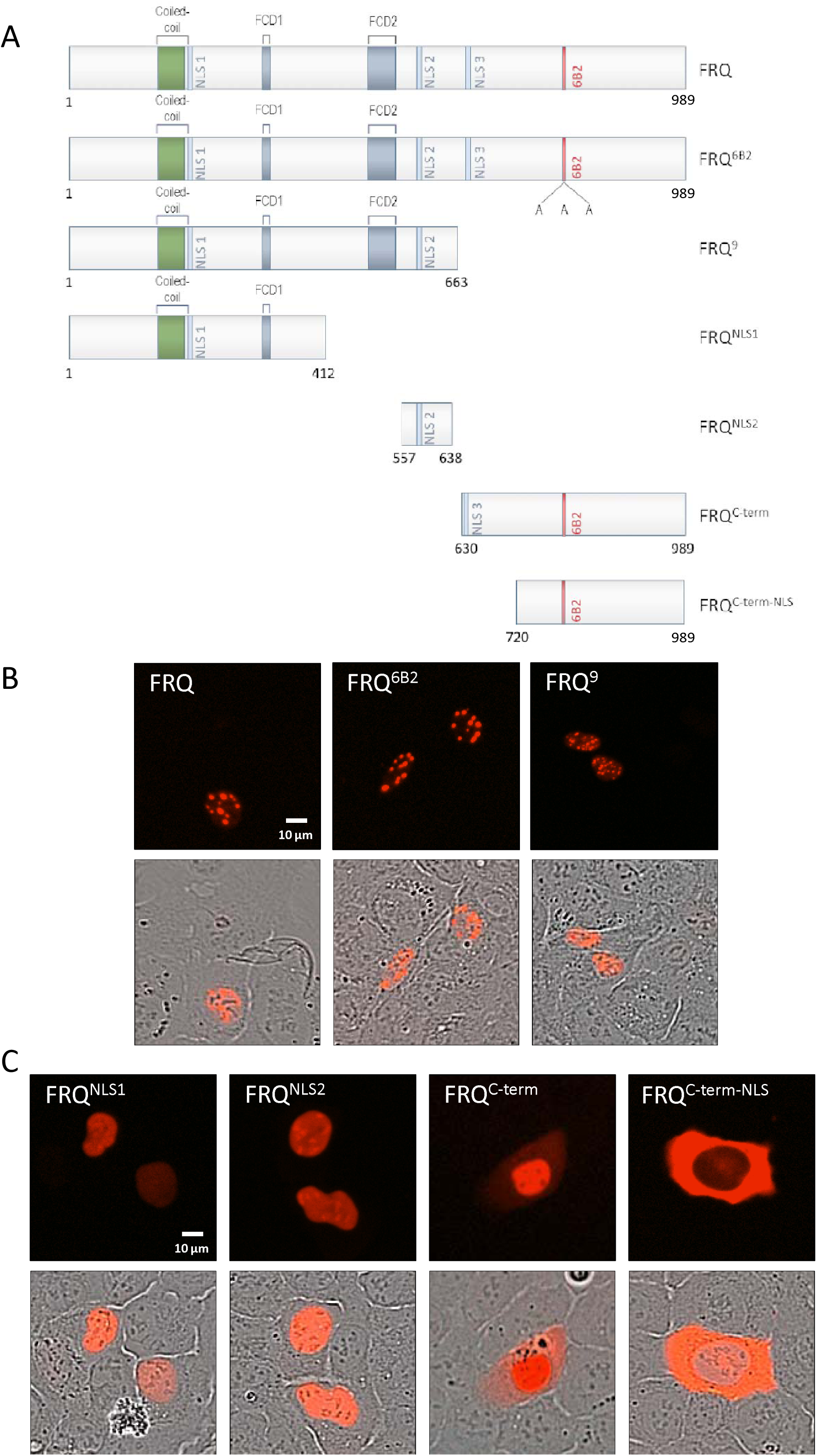
FRQ contains three NLSs. **A** Schematic of FRQ constructs. The coiled-coil dimerization domain, the predicted NLSs, the FRQ-CK1a interaction domains (FCD) 1 and 2, as well as the 6B2 region required for FRH binding ^7,8,21^ are indicated. The FRQ^6B2^ mutant carries an alanyl substitution of residues 774-776, DHF to AAA, which are part of a larger binding site ^21,22^. The proteins were tagged either N-terminally with mK2 or C-terminally with mNG. **B, C** Transient expression in U2OStx cells of the indicated mK2-tagged FRQ versions.

FRQ contains three polybasic sequences that could function as nuclear localization signal (NLS) (Fig. 1A, S1C). We expressed mK2-tagged fragments of FRQ, each containing one of the three polybasic sequences, mK2-FRQ^NLS1^ (aa1-412), mK2-FRQ^NLS2^ (aa557-638), and mK2-FRQ^C-term^ (aa630-989) (Fig. 1A). The proteins localized to the nucleus, although a substantial portion of mK2-FRQ^C-term^ was detected in the cytosol. In contrast, mK2-FRQ^C-term-NLS^ (aa720-989), a C-terminal fragment lacking the polybasic sequence, accumulated in the cytosol (Fig. 1A, C). These data suggest that FRQ contains three NLSs, henceforth referred to as NLS1, NLS2, and NLS3.

We noticed that the smaller FRQ fragments showed a significantly reduced tendency to form foci. This suggests that the entire FRQ molecule contributes to foci formation, presumably through self-interaction. Recombinant FRQ has been recently reported to phase-separate *in vitro*, and focal assemblies of FRQ have been described in *Neurospora* ^24^. It will be interesting to analyze if FRQ’s potential to self-interact has functional significance.

NLS1 has previously been identified and a strain expressing FRQ^ΔNLS1^ was reported to be arrhythmic ^25^. However, NLS1 is adjacent to the coiled-coil region that mediates dimerization of FRQ, and deletion of NLS1 interferes with FRQ dimerization ^26^, which is essential for its function ^3^.

We therefore generated a mK2-FRQ versions where the three NLSs were mutated rather than deleted (Fig. S1D). mK2-FRQ^mNLS1/2/3^ localized to the cytoplasm of U2OStx cells (Fig. S1E, left panel), indicating FRQ does not contain additional NLSs.

### Impact of NLSs on FRQ function in *Neurospora*

We then created *Neurospora* strains expressing FRQ versions with mutated NLSs under control of the native *frq* promoter, alongside a control strain expressing wild-type (WT) FRQ. These strains also express a *frq-lucP* reporter to enable analysis of circadian rhythms *in vivo* through bioluminescence recordings. The strains, grown in 96-well plates, were exposed to 12 h light, 12 h dark and 12 h light and then transferred to the dark. The control strain, expressing WT FRQ, displayed a robust bioluminescence rhythm in constant darkness (Fig. 2A). Strains expressing FRQ^mNLS1^ (NLS1: RRKKR to RQKKQ) displayed a WT period which slowly damped (Fig. 2B), indicating that FRQ^mNLS1^ is a dimer in contrast to FRQ^ΔNLS1^, which has lost all clock functions ^26^. This finding suggests that the arrhythmic phenotype of FRQ^ΔNLS1^ is likely due to the disrupted FRQ dimerization.

**Fig. 2.**
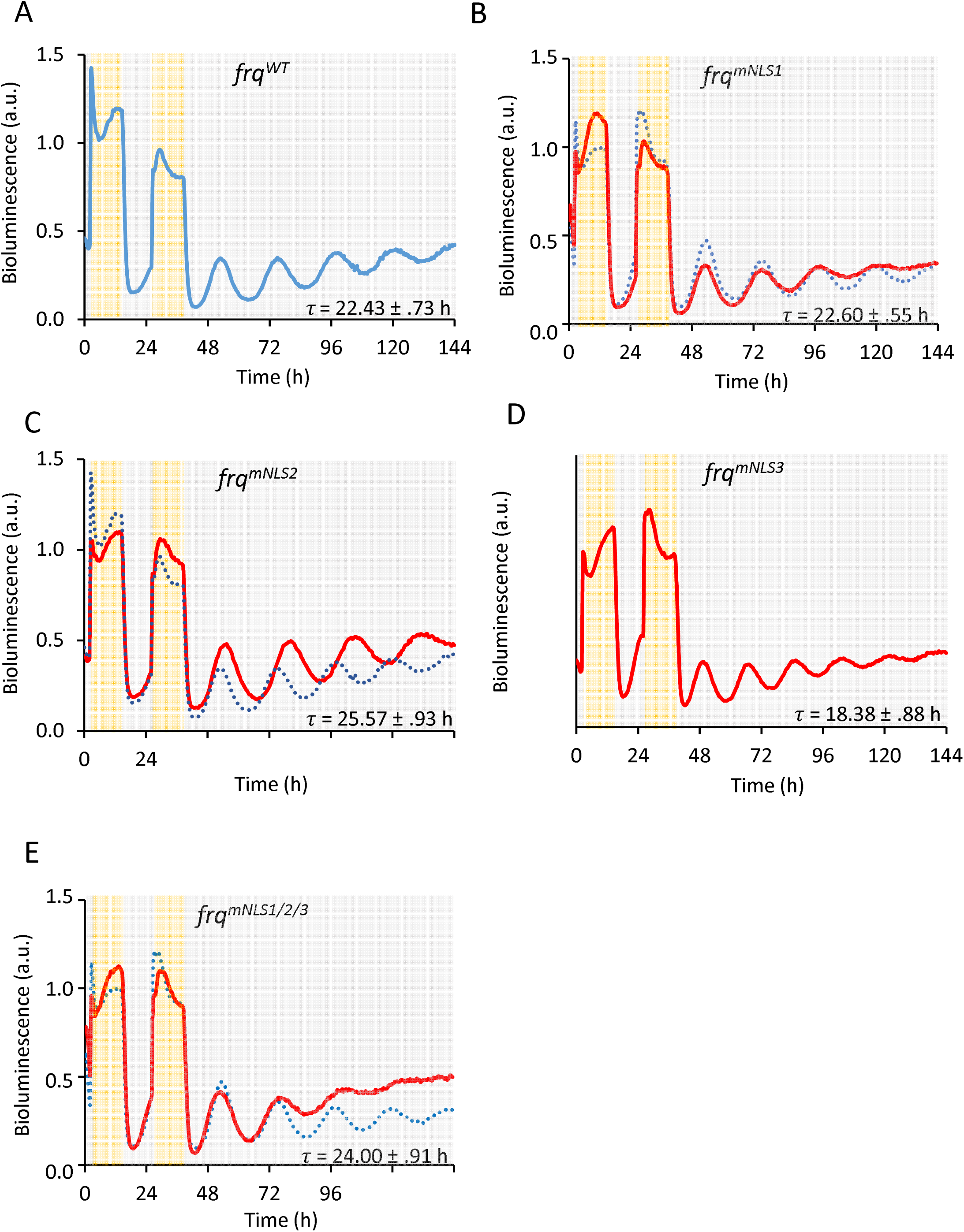
Impact of FRQ’s NLSs on the circadian clock. Bioluminescence recordings of *Neurospora frq-lucP* reporter strains expressing the indicated *frq* alleles. Calculated period lengths (±SD) are shown. **A** WT *frq*. The trajectory is the normalized average of 12 replicates from 2 independent clones. Also shown as dotted blue line for comparison in B-E. **B** *frq^mNL1^*. 16 replicates, 4 clones. **C** *frq^mNL2^*. 12 replicates, 3 clones. **D** *frq^mNLS3^*. 12 replicates, 3 clones. **E** *frq^mNLS1/2/3^*. 8 replicates, 2 clones.

Strains expressing FRQ^mNLS2^ (NLS2: RRKKRK to RGQERK) displayed a long period rhythm and strains expressing FRQ^mNLS3^ (NLS3: RRKRR to RGQGR) displayed an early circadian phase and a short period rhythm dampening slightly after several days in the dark (Fig. 2C, D). A triple mutant, FRQ^mNLS1/2/3^, displayed a low amplitude rhythm that damped rapidly in constant darkness (Fig. 2E). The data indicate that the NLSs are crucial for robust circadian oscillations, although some residual clock functions are retained in their absence, suggesting that FRQ^mNLS1/2/3^ must have additional means to enter nuclei in *Neurospora*, potentially through interaction with FRH.

### FRH interferes with FRQ self-interaction

FRQ forms a tight complex with FRH, which protects FRQ from premature degradation in *Neurospora* ^4^. We constructed a plasmid encoding FRH C-terminally tagged with mNG and expressed the protein in U2OStx cells. FRH-mNG was evenly distributed in the nucleus (Fig. S1F, G). When co-expressed with FRQ, the phenotype depended on the ratio and amounts of the transfected expression vectors (Fig. 3A, B). At a plasmid DNA transfection ratio of 16.5 ng FRH-mNG to 50 ng mK2-FRQ (Fig. 3A), FRH-mNG co-localized with mK2-FRQ in nuclear foci. However, at a 50 ng : 50 ng transfection ratio, both proteins were evenly distributed in the cell nucleus (Fig. 3B). These data indicate that the proteins interacted at both expression ratios, but higher levels of FRH-mNG impaired the accumulation of mK2-FRQ in nuclear foci. This suggests that FRH-mNG prevented the self-interaction of mK2-FRQ.

**Fig. 3.**
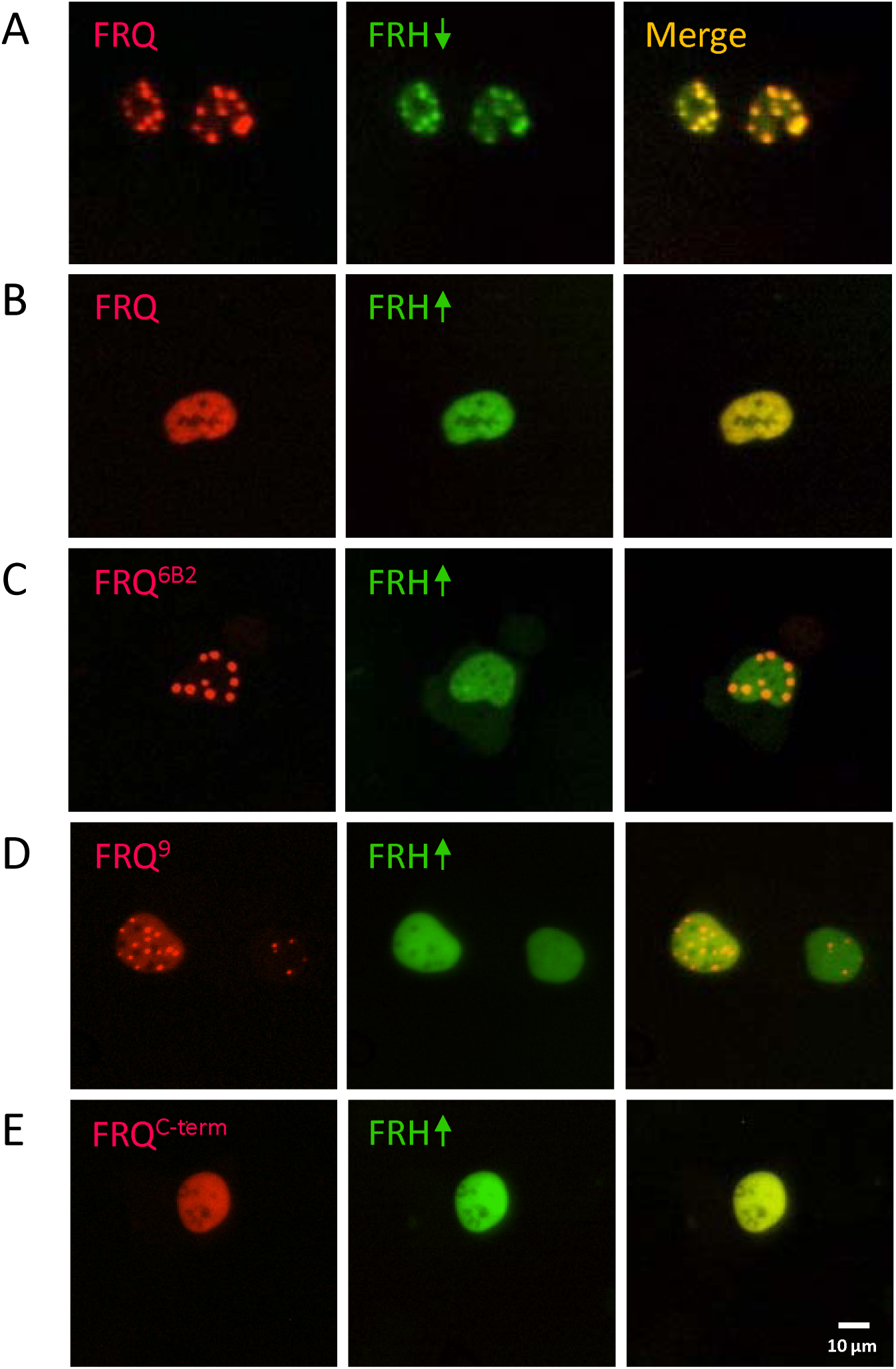
Binding of saturating amounts of FRH to FRQ suppresses the formation of FRQ nuclear foci in U2OStx cells. The indicated mK2-tagged FRQ versions were co-expressed with FRH-mNG. **A** Co-expression of mK2-FRQ (50 ng plasmid DNA) and non-saturating amounts of FRH-mNG (16.5 ng). mK2-FRQ and FRH-mNG co-localize in nuclear foci. **B** mK2-FRQ (50 ng) and saturating amounts of FRH-mNG (50 ng). mK2-FRQ and FRH-mNG are dispersed in the nucleus. **C, D** 50 ng mK2-FRQ^6B2^ or mK2-FRQ^9^ together with FRH-mNG (50 ng). FRH-mNG is dispersed in the nucleus while C mK2-FRQ^6B2^ and **D** mK2-FRQ^9^ form nuclear foci. **E** mK2-FRQ^C-term^ (50 ng) and FRH-mNG (50 ng). Both proteins are dispersed in the nucleus.

We then expressed mK2-FRQ^6B2^ with FRH-mNG. mK2-FRQ^6B2^ accumulated in nuclear foci, and these foci did not contain FRH-mNG, which was homogeneously dispersed in the nucleus (Fig. 3C). Similarly, the C-terminally truncated protein mK2-FRQ^9^ accumulated in nuclear foci, which was not affected by FRH-mNG (Fig. 3D).

The data suggest that the tight anchoring of FRH to its binding site in the 6B2 region of FRQ prevented the accumulation of FRQ in nuclear foci, a process requiring regions of FRQ outside the FRH anchoring site. In the absence of site-specific tight anchoring, FRH does not interact with FRQ. Thus, anchoring FRH increases its local concentration, facilitating thermodynamically weak interactions with additional parts of FRQ that would otherwise remain undetectable. Such interactions of FRH with multiple regions distributed across the entire FRQ have been recently proposed on the basis of peptide array studies ^22^. In this way, anchored FRH effectively competes with FRQ-FRQ self-interactions.

We then co-expressed mK2-FRQ^C-term^ with FRH-mNG (Fig. 3E). As expected from the localization of the individually expressed proteins, both proteins were evenly distributed throughout the nucleus. However, upon comparing data shown in Fig. 1C and 3E, we observed that the nuclear localization of mK2-FRQ^C-term^ was more pronounced when co-expressed with FRH. These findings suggest that FRH enhances the nuclear localization of mK2-FRQ^C-term^.

We therefore co-expressed mK2-FRQ^mNLS1/2/3^ with FRH-mNG. mK2-FRQ^mNLS1/2/3^ was recruited into the nucleus (Fig. S1E, right panel), indicating that FRH can drive import of FRQ via its own NLS.

### FRQ interacts with the WCC in absence of FRH

In *Neurospora*, WC-1 is rapidly degraded without its stabilizing partner WC-2 and, hence, cannot be easily analyzed. We tagged WC-1 with mNG and WC-2 with mK2 to study their expression and localization in U2OStx cells. WC-1-mNG was well expressed and localized predominantly in the cytosol (Fig. S2A), while mK2-WC-2 localized to the nucleus and formed droplet-like foci (Fig. S2B). When expressed together, WC-1-mNG was depleted from the cytosol and co-localized with mK2-WC2 in nuclear foci (Fig. S2C), indicating interaction of the two WCC subunits.

In *Neurospora*, the FFC complex interacts weakly and dynamically with the WCC ^4,9^ and inactivates WCC through phosphorylation by CK1a ^9^.

To investigate whether FRH is necessary for this interaction, we first co-expressed in U2OStx cells WC-1-mNG with mK2-FRQ in absence of WC-2 and FRH. WC-1-mNG, which localized to the cytosol when expressed by itself (Fig. S2A), co-localized completely with mK2-FRQ in nuclear foci (Fig. 4A), demonstrating that WC-1 by itself interacted with FRQ. Upon co-expression with mK2-FRQ^6B2^, which has a mutation in the FRH binding site, WC-1-mNG was also recruited into the nucleus, but a substantial fraction of WC-1-mNG remained cytosolic (Fig. 4B). The data suggest that WC-1-mNG interacted with mK2-FRQ^6B2^ but potentially with reduced affinity. In contrast, WC-1-mNG remained in the cytosol when co-expressed with the C-terminally truncated mK2-FRQ^9^, which formed nuclear foci that did not contain WC-1-mNG (Fig. 4C). Finally, WC-1-mNG was recruited and uniformly dispersed in the nucleus when co-expressed with mK2-FRQ^C-term^ (Fig. 4D). These findings indicate that the C-terminal third of FRQ is necessary and sufficient for interaction with WC-1, while the co-localization assay did not detect an interaction between WC-1-mNG and mK2-FRQ^9^, corresponding to the N-terminal two-thirds of FRQ.

**Fig. 4:**
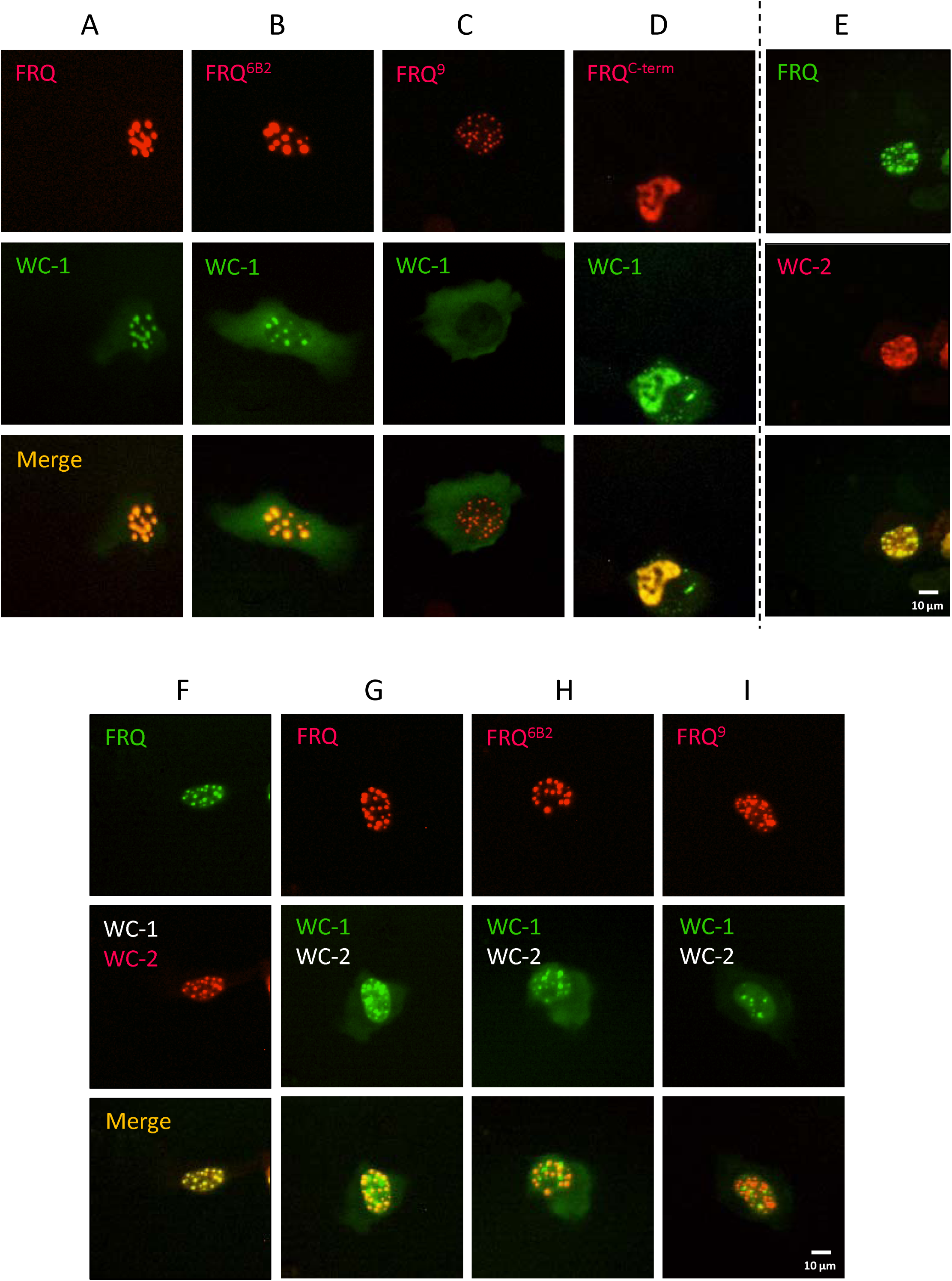
FRQ interacts with WC1 and WC-2 in absence of FRH. **A** Co-localization of mK2-FRQ and WC-1-mNG in nuclear foci. **B** mK2-FRQ^6B2^ and WC-1-mNG co-localize in nuclear foci. **C** mK2-FRQ^9^ forms nuclear foci while co-expressed WC-1-mNG remains in the cytosol. **D** mK2-FRQ^C-term^ is homogeneously dispersed in the nucleus and recruits WC-1-mNG from the cytosol to the nucleus. **E** FRQ-mNG and mK2-WC-2 co-localize in nuclear foci. **F** Co-expression of FRQ-mNG with untagged WC-1 and mK2-WC-2. FRQ-mNG and mK2-WC-2 co-localize in nuclear foci. **G** Co-expression of mK2-FRQ with WC-1-mNG and untagged WC-2. mK2-FRQ and WC-1-mNG co-localize in nuclear foci. **H** Expression of mK2-FRQ^6B2^ with WC-1-mNG and untagged WC-2. mK2-FRQ^6B2^ and WC-1-mNG co-localize in nuclear foci. **I** Expression of mK2-FRQ^9^ with WC-1-mNG and untagged WC-2. FRQ^9^-mNG and mK2-WC-2 accumulate in distinct, nonoverlapping nuclear foci.

Next, we expressed mK2-WC-2 together with FRQ-mNG (Fig. 4E). Both proteins formed nuclear foci, which overlapped, suggesting that WC-2 on its own also interacts with FRQ. To study the interaction of FRQ with the WCC, we co-expressed FRQ-mNG with untagged WC-1 and mK2-WC-2 (Fig. 4F) as well as mK2-FRQ with WC-1-mNG and untagged WC-2 (Fig. 4G). As expected, FRQ-mNG as well as mK2-FRQ formed nuclear foci. The FRQ-mNG foci overlapped with mK2-WC-2 foci (Fig. 4F) and similarly, mK2-FRQ foci overlapped with WC-1-mNG (Fig. 4G), suggesting interaction of FRQ with WCC in absence of FRH. Some WCC foci remained separate, potentially due to the saturation of the FRQ foci with WCC, causing excess WCC to accumulate in distinct foci.

When WC-1-mNG and untagged WC-2 were co-expressed with mK2-FRQ^6B2^ the foci overlapped (Fig. 4H), indicating that the 6B2 region is not required for the interaction of FRQ with WCC. A few WC-1-mNG (WCC) foci remained distinct, suggesting excess of WCC over mK2-FRQ^6B2^. In contrast, when WC-1-mNG and untagged WC-2 were co-expressed with mK2-FRQ^9^, foci of WC-1-mNG (WCC) and mK2-FRQ^9^ did not overlap (Fig. 4I), demonstrating that the proteins did not interact.

Recent deletion studies suggested that the N-terminal half of FRQ is required for its interaction with WCC in *Neurospora* ^27^. However, as these deletions are predicted to disrupt FRQ dimerization (e.g. FRQ^Δ149–193^) and CK1a recruitment (e.g. FRQ^Δ482-510^), they may impact WCC interaction indirectly.

The overexpression and co-localization in U2OStx cells indicate that the C-terminal part of FRQ is necessary and sufficient to interact with both WC-1 and WC-2, even in the absence of FRH. Because the 6B2 mutation in FRQ weakens but does not eliminate WCC binding, it suggests that the WCC binding region overlaps with, but is distinct from, the FRH binding site. This raises the question of how FRH modulates this interaction.

### FRH blocks WCC binding to FRQ

WCC and FRH both bind to the C-terminal part of FRQ. Since WC-1-mNG is cytoplasmic on its own but recruited into the nucleus by mK2-FRQ, this allows for easy quantification of the impact of FRH on this interaction. We therefore co-expressed WC-1-mNG and mK2-FRQ^C-term^ with and without untagged FRH. When WC-1-mNG was co-expressed with mK2-FRQ^C-term^ it was enriched in the nucleus and was homogeneously dispersed just like mK2-FRQ^C-term^ (Fig. 5A, left). When untagged FRH was also co-expressed, mK2-FRQ^C-term^ displayed nuclear localization, while WC-1-mNG remained predominantly in the cytoplasm (Fig. 5A, right), just like in the absence of mK2-FRQ^C-term^ (see Fig. S2A). Similarly, when WC-1-mNG and mK2-FRQ were co-expressed, WC-1-mNG was recruited from the cytosol into the nucleus and colocalized with mK2-FRQ in nuclear foci (Fig. 5B, left). In presence of FRH, mK2-FRQ was homogeneously dispersed in the nucleus, indicating its interaction with saturating amounts of FRH, while WC-1-mNG remained predominantly cytoplasmic (Fig. 5B, right).

**Fig. 5:**
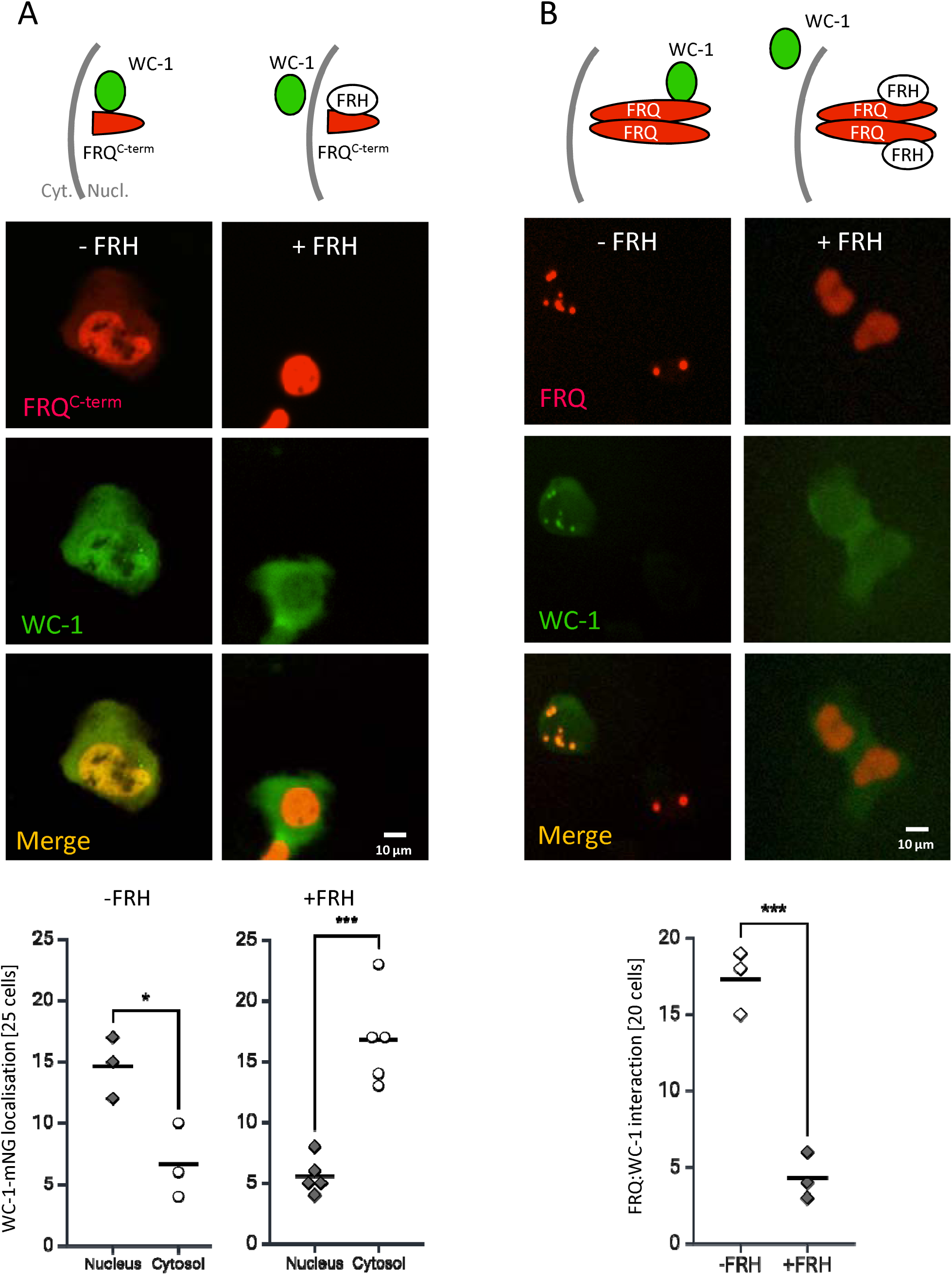
FRH interferes with binding of FRQ to WC-1. Schematics depict the interactions of fluorescently tagged (green and red) and untagged (white) proteins. **A** Co-expression of mK2-FRQ^C-term^ and WC-1-mNG without FRH and with FRH. FRH prevents the recruitment of WC-1-mNG into the nucleus. 25 cells per experiment were analyzed. - FRH: 3 independent experiments, + FRH: 5 independent experiments. Unpaired T-test, *: p ≤ 0.05; ***: p ≤ 0.001. **B** Co-expression of mK2-FRQ and WC-1-mNG without FRH and with FRH. Interaction of mK2-FRQ with WC-1-mNG was evaluated by counting cells displaying co-localization in nuclear foci (-FRH) or cells displaying nuclear enrichment of WC-1-mNG (+FRH), which is significantly reduced in presence of FRH. 20 cells each from three independent experiments were evaluated. Unpaired T-test, ***: p ≤ 0.001.

To further analyze the impact of FRH on the interaction of FRQ with the WCC, we co-expressed WC-1-mNG and untagged WC-2 together with mK2-FRQ and either low amounts of untagged FRH, allowing co-localization of FRQ and FRH in nuclear foci, or with high amounts of FRH, interfering with foci formation (see Fig. 3A and B). At low levels of FRH (Fig. S3A, left), mK2-FRQ accumulated in nuclear foci. WC-1-mNG co-localized with these foci, suggesting WCC interacted with FRQ. At high levels of FRH (Fig. S3A, right), mK2-FRQ was homogeneously dispersed in the nucleus, indicating that saturating amounts of FRH interfered with the self-interaction of mK2-FRQ. WC-1-mNG accumulated in nuclear foci, indicating that it assembled with untagged WC-2. The WC-1-mNG (WCC) foci, however, did not contain mK2-FRQ (Fig. S3A, right). The data suggest that FRQ saturated with FRH does not interact with WCC.

For additional analysis, we expressed mK2-WC-2 and untagged WC-1 together with untagged FRQ and either high or low amounts of FRH-mNG. At low amounts, FRH-mNG was enriched in mK2-WC-2 (WCC) foci (Fig. S3B, left), indicating that unlabeled FRQ in complex with subsaturating amounts of FRH-mNG interacted with WCC. When expressed at a high level, FRH-mNG was dispersed in the nucleus, while mK2-WC-2 (WCC) foci persisted (Fig. S3B, right).

The data strongly suggest that FRH is not only unnecessary for the interaction of FRQ with WC-1 and WC-2, but it may even interfere with it. WCC and FRH bind to overlapping but distinct sites within the C-terminal region of FRQ. The strong anchoring of FRH inhibits WCC binding when FRH is present in high, presumably saturating amounts, but permits WCC-FRQ interaction at lower, subsaturating FRH concentrations. This finding was surprising and unexpected, as it had been suggested that FRH is necessary for the interaction between WCC and FFC ^28^. This suggestion was based on data showing convincingly that WCC did not interact with FRQ complexes containing a mutant FRH with an R806H substitution.

To investigate further, we expressed WC-1-mNG and mK2-FRQ with either unlabeled FRH or FRH^R806H^. Both FRH and FRH^R806H^ competed with the nuclear recruitment of WC-1-mNG by mK2-FRQ (Fig. S4). However, FRH^R806H^ was a stronger competitor than FRH, supporting the notion that the R806H substitution acts as a dominant-negative mutation, leading to tighter binding to FRQ, potentially explaining why FRQ did not interact with WCC in the *frh^R806H^ Neurospora* strain ^28^.

### Phosphorylation of FRQ by CK1 induces its dissociation from FRH and nuclear export

Having shown that FRH blocks binding of WCC to FRQ, we asked whether and how the interaction of FRQ with WCC is regulated. In *Neurospora*, FRQ is gradually hyperphosphorylated by CK1a, which inactivates FRQ and favors its degradation ^10–12^. To analyze the impact of CK1a, we co-expressed in U2OStx cells untagged CK1a with FRH-mNG and either mK2-FRQ, mK2-FRQ^6B2^ or mK2-FRQ^9^. The three FRQ proteins were hyperphosphorylated in a CK1a-dependent fashion (Fig. 6A). This indicates that CK1a phosphorylated mK2-FRQ in U2OStx cells while endogenous kinases did not support detectable hyperphosphorylation of the overexpressed FRQ proteins.

**Fig. 6:**
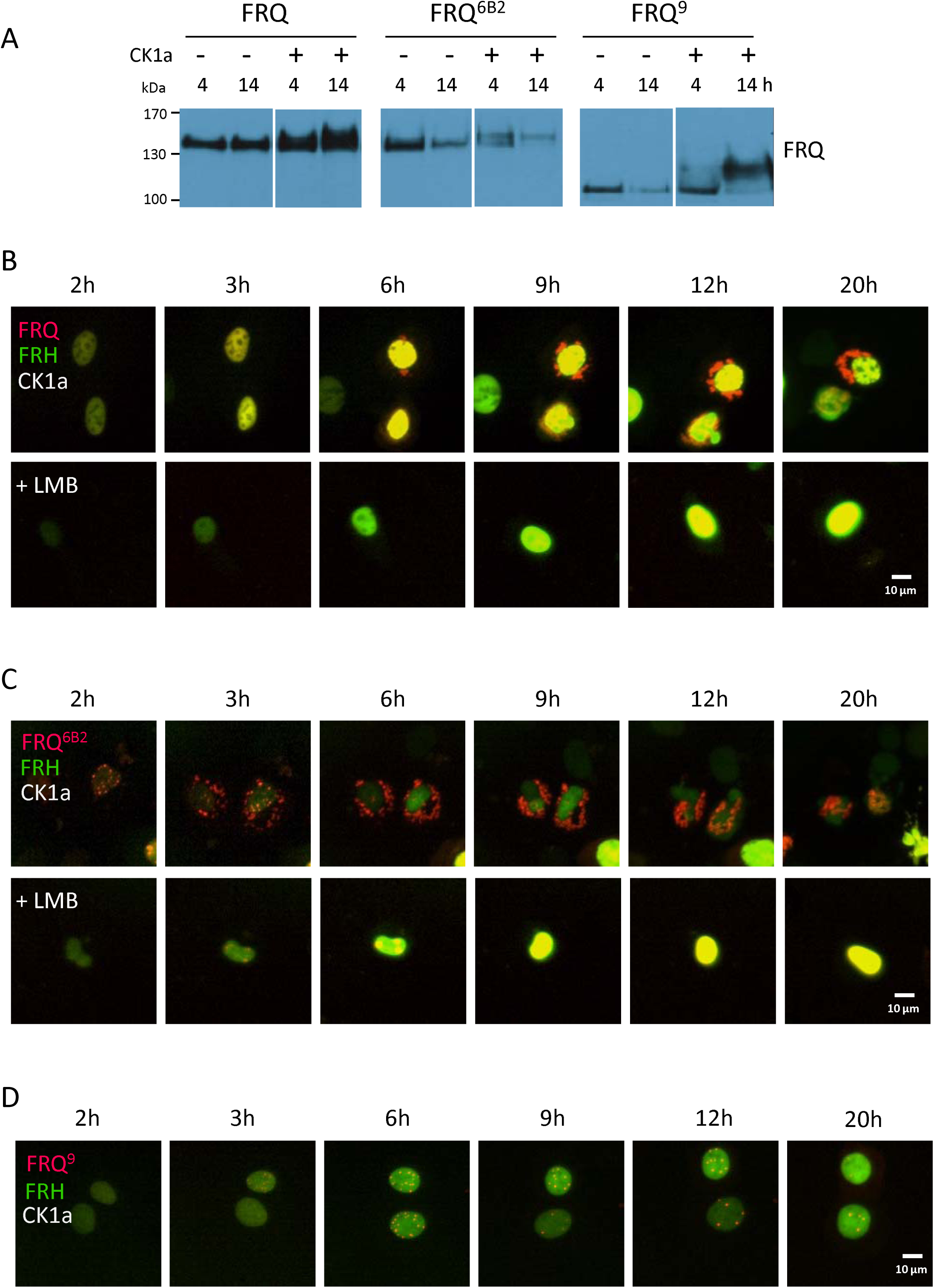
Phosphorylation of FRQ by CK1a triggers its dissociation from FRH and nuclear export. **A** Western blot analysis of mK2-FRQ, mK2-FRQ^6B2^ and mK2-FRQ^9^ co-expressed with and without CK1a in U2OStx cells for the indicated time periods. The samples were loaded side by side on the same gel in the order shown. They were cropped into separate panels in pairs to facilitate comparison. **B - D** Incucyte time courses of transfected U2OStx cells upon co-expression of FRH-mNG and untagged CK1a with **B** mK2-FRQ, **C** mK2-FRQ^6B2^ and, **D** mK2-FRQ^9^. LMB was added when indicated.

We then analyzed the dynamics of CK1a’s effect on mK2-FRQ and FRH-mNG by live-cell microscopy. Initially, mK2-FRQ and FRH-mNG co-localized in the nucleus. mK2-FRQ did not form foci, indicating that it interacted with saturation levels of FRH (Fig. 6B). After several hours, CK1a induced the formation of amorphous assemblies of mK2-FRQ in the cytosol, while FRH-mNG remained confined to the nucleus (Fig. 6B, S5A). When CRM-1-dependent nuclear export was inhibited with leptomycin B, mK2-FRQ remained in the nucleus (Fig. 6B lower panels, S5A). These data indicate that phosphorylation by CK1a caused dissociation of mK2-FRQ from FRH-mNG and supported its export from the nucleus. mK2-FRQ^6B2^, which cannot interact with FRH-mNG, was also exported from the nucleus in an LMB-sensitive manner when co-expressed with FRH-mNG and CK1a, but with substantially faster kinetics than mK2-FRQ (Fig. 6C, S5B), indicating that FRQ-bound FRH delayed nuclear export. The data suggest that phosphorylation inactivated the positively charged NLSs, potentially through neutralization by negatively charged phosphosites or -clusters generated by CK1a. In contrast, mK2-FRQ^9^ was not exported from the nucleus when co-expressed with CK1a and FRH-mNG (Fig. 6D, S5C), suggesting that the C-terminal part of FRQ contains a nuclear export signal (NES).

To further narrow down the NES, we divided the C-terminus of FRQ into three parts, deleting amino acid residues 631-756, 757-888, and 889-989 in mK2-FRQ (Fig. S6A), respectively, and expressed each modified protein along with CK1a in U2OStx cells (Fig. S6B). After DOX induction, mK2-FRQ^Δ631-756^ and mK2-FRQ^Δ889-989^ accumulated to the cytosol, suggesting that CK1a-mediated phosphorylation favored cytoplasmic localization by inactivating the NLSs and/or activating the NES. In contrast, mK2-FRQ^Δ757-888^ remained nuclear, demonstrating its NES had been deleted. Since this deleted segment also includes the FRH binding region, it seems likely that FRH can mask the NES, thereby regulating FRQ’s nuclear export in a phosphorylation-dependent manner.

Our data indicate that in the absence of phosphorylation, the three NLSs in full-length FRQ dominate functionally over the NES, resulting in strong nuclear enrichment of FRQ. When FRQ is hyperphosphorylated by CK1a, it is exported to the cytosol, likely due to NLS neutralization. FRH slows this phosphorylation-driven export of FRQ by providing additional nuclear import capacity via its NLS and by masking FRQ’s NES.

### Native FFC dissociates upon phosphorylation of FRQ by CK1a

In *Neurospora*, all detectable FRQ is present in a complex with FRH, which protects FRQ from degradation. FRQ^6B2^ or FRQ^9^, which cannot interact with FRH, are rapidly degraded. Hence, they accumulate only at low levels, despite being synthesized in excessive amounts due to nonfunctional negative feedback ^21,29^.

Since free FRQ does not accumulate at significant levels in *Neurospora*, it is challenging to investigate whether gradual hyperphosphorylation triggers its dissociation from FRH. Proteasome inhibitors are not very effective and lead to pleiotropic effects when administered at high doses and for longer time periods. To circumvent degradation of free FRQ, we prepared native *Neurospora* extract from cells expressing 2xFLAG-tagged FRH and hyperphosphorylated FRQ *in vitro* by adding recombinant CK1a and ATP (Fig. 7A). An untreated extract was used for comparison. Subsequently, 2xFLAG-FRH was subjected to immunoprecipitation. In the control reaction, hypophosphorylated FRQ co-immunoprecipitated with 2xFLAG-FRH (Fig. 7B left). In contrast, upon incubation with CK1a and ATP, hyperphosphorylated FRQ no longer co-immunoprecipitated with 2xFLAG-FRH (Fig. 7B right).

**Fig. 7:**
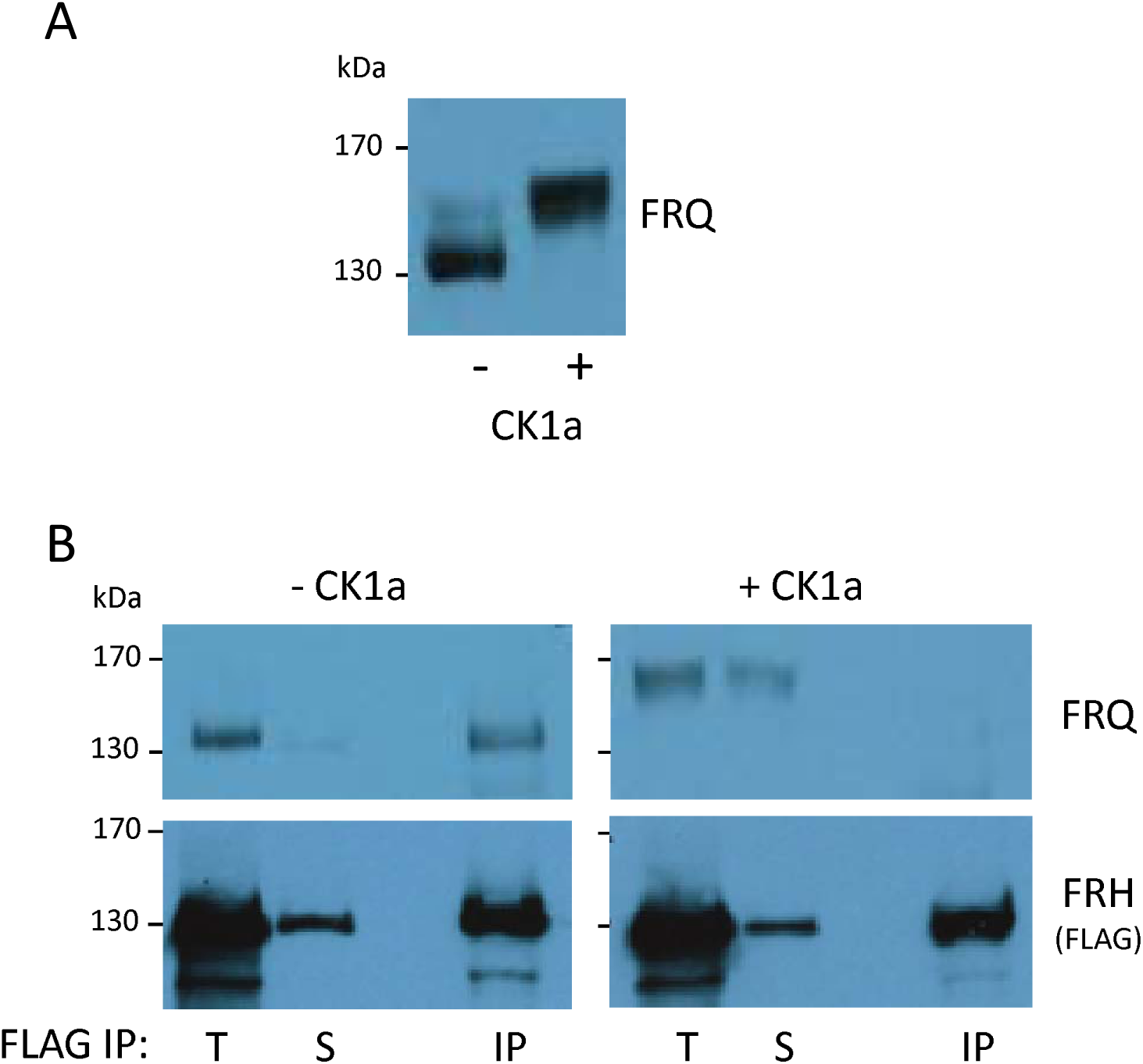
Phosphorylation of native FRQ by CK1a triggers its dissociation from FRH. **A** Whole cell lysate (WCL) of *Neurospora* expressing 2xFLAG-tagged FRH prepared from cultures containing hypophosphorylated FRQ (grown 3 h in light after 10 h darkness) was incubated with recombinant CK1a and ATP, or left untreated. **B** Lysates from **A** were subjected to FLAG immunoprecipitation. Total (T), supernatant (S), and immunoprecipitate (IP) were analyzed by Western blot with antibodies against FRQ (upper panels and 2xFLAG-FRH (Lower panels). n = 3.

Size exclusion chromatography of untreated native *Neurospora* cell extract revealed that FRH co-eluted with FRQ in a high molecular mass fraction, indicating their interaction (Fig. S7A, B). Given FRH’s abundance relative to FRQ, most FRH molecules eluted in a lower molecular mass fraction, consistent with prior findings ^26^. Conversely, when the cell extract was treated with CK1a and ATP, FRH no longer co-eluted with FRQ in the high molecular mass fraction (Fig. S7A, B).

These findings strongly suggest that the phosphorylation of native FRQ by CK1a triggers its dissociation from FRH.

## Discussion

We have previously shown that CK1a, anchored to FRQ, slowly hyperphosphorylates FRQ in a temperature-independent manner, forming a molecular module suitable for time measurement on a circadian scale ^16^. However, the molecular mechanism underlying phosphorylation-dependent circadian timekeeping remains unclear. In this study, we provide data demonstrating that FRH, the folded binding partner of the intrinsically disordered FRQ, senses the time-dependent phosphorylation state of FRQ, forming an elaborate molecular timing device. The key features of this timer rely on a series of previously unanticipated competitive interactions regulated by slow phosphorylation of FRQ by bound CK1a. We demonstrate that the slow gradual phosphorylation of FRQ initiates a two-step subunit remodeling process, first activating and then inactivation the FRQ-FRH complex in a time-dependent manner. Our data identify FRH as a decoding hub for temporal information encoded on FRQ through accumulated phosphate groups.

### FRH anchored to the C-terminal part of FRQ interacts with additional parts of FRQ

Using live-cell microscopy, we visualized the interactions and dynamics of *Neurospora* clock proteins in U2OStx cells. This heterologous cell-based approach allowed us not only to characterize clock proteins but also to analyze their interactions and subcellular dynamics, which could not be examined in *Neurospora* or in a cell-free system with purified components. In U2OStx cells, overexpressed FRQ accumulated in nuclear foci, suggesting a propensity for self-interaction. Interestingly, co-expression of high levels of FRH prevented the formation of nuclear FRQ foci but did not prevent foci formation by FRQ^9^ (aa 1–662) or FRQ^6B2^, which both lack the FRH anchor site. These data suggest that anchoring FRH to the 6B2 region of FRQ increases the local concentration of FRH, favoring its weaker interaction with other regions of FRQ over FRQ self-interactions. The nuclear foci formed by overexpressed FRQ in U2OS cells may represent phase-separated droplets. In *Neurospora*, FRQ is expressed at lower levels but may also accumulate in larger assemblies ^24^. It remains to be determined whether the self-interaction of FRQ and its suppression by FRH occur only at the level of a FRQ dimer or also in higher oligomeric assemblies.

### FRH dissociates from phosphorylated FRQ

Our most surprising finding was that, following a time delay, phosphorylation of FRQ by CK1a triggered the dissociation of the FRQ-FRH complex, resulting in the nuclear export of FRQ, while FRH remained in the nucleus of U2OStx cells. This was unexpected, as unbound FRQ is inherently unstable and rapidly degraded in *Neurospora*, which has hindered detailed analyses of its subcellular dynamics in absence of FRH. We further validated this phosphorylation-dependent dissociation by analyzing native FRQ-FRH complexes from *Neurospora* through co-immunoprecipitation and by size-exclusion chromatography.

### FRQ subcellular localization is regulated by three NLSs and one NES and by FRH

We found that FRQ contains three NLSs in the N-terminal two-thirds and one CRM1-dependent NES in the C-terminal third overlapping the FRH binding site. The NLSs are functionally dominant over the NES, leading to the nuclear accumulation of FRQ, including FRQ^6B2^, which cannot bind FRH. This nuclear accumulation is further enhanced by FRH, which has its own NLS and masks the NES when bound to FRQ. Phosphorylation of FRQ by CK1a promotes its nuclear export, likely by inhibiting the polybasic NLSs through interactions with phosphorylated regions in FRQ and by facilitating the release of bound FRH.

### FRH masks the WCC binding-site and is released upon phosphorylation of FRQ

It has been reported that specific mutations in the C-terminal third of FRQ affect its interaction with WCC but not with FRH ^27^. We show here that FRQ interacts with both WCC subunits, WC-1 and with WC-2. The C-terminal part of FRQ is necessary and sufficient for the interaction with WCC. Surprisingly, we found that the associations of FRQ with WCC and FRH are mutually exclusive. The binding sites in FRQ are overlapping but not identical, and the tight anchoring of FRH to unphosphorylated FRQ blocks the WCC binding site. This site becomes accessible only when FRH is released through FRQ phosphorylation by CK1a.

Our data suggest fundamental modifications of the *Neurospora* circadian clock model. Specifically, we show that the phosphorylation state of the intrinsically disordered FRQ dimer is decoded by its folded partner, FRH. Slow multisite phosphorylation of FRQ by CK1a triggers with a delay remodeling of an initially inactive nuclear hetero-tetrameric complex with FRQ_2_FRH_2_ stoichiometry (and bound CK1a) into an active trimeric FRQ_2_FRH_1_ complex that interacts with and inhibits WCC. Phosphorylation also inactivates FRQ’s NLSs. The trimeric FRQ_2_FRH_1_ complex may shuttle between compartments, exported via the NES exposed by release of FRH from one FRQ subunit and reimported as long as the six NLSs of the FRQ dimer are not fully inactivated by phosphorylation and in addition via FRH bound to the second FRQ subunit. Because FRQ phosphorylation is inherently slow and at least partially stochastic, the two FRH molecules are not released simultaneously. Instead, the second FRH is released only after a delay, leading to the formation of the FRQ_2_ complex. In the nucleus, the FRQ_2_ complex is active in negative feedback regulation. However, once exported to the cytosol, phosphorylated FRQ_2_ cannot be reimported and is eventually degraded. This stepwise subunit remodeling of the FFC converts the steadily increasing phosphorylation state of FRQ into two distinct molecular switches: a delayed ‘on-switch’ that activates previously accumulated FRQ by exposing its WCC binding site, and an ‘off-switch’ that inactivates FRQ by promoting its nuclear export and degradation (Fig. 8).

**Fig. 8:**
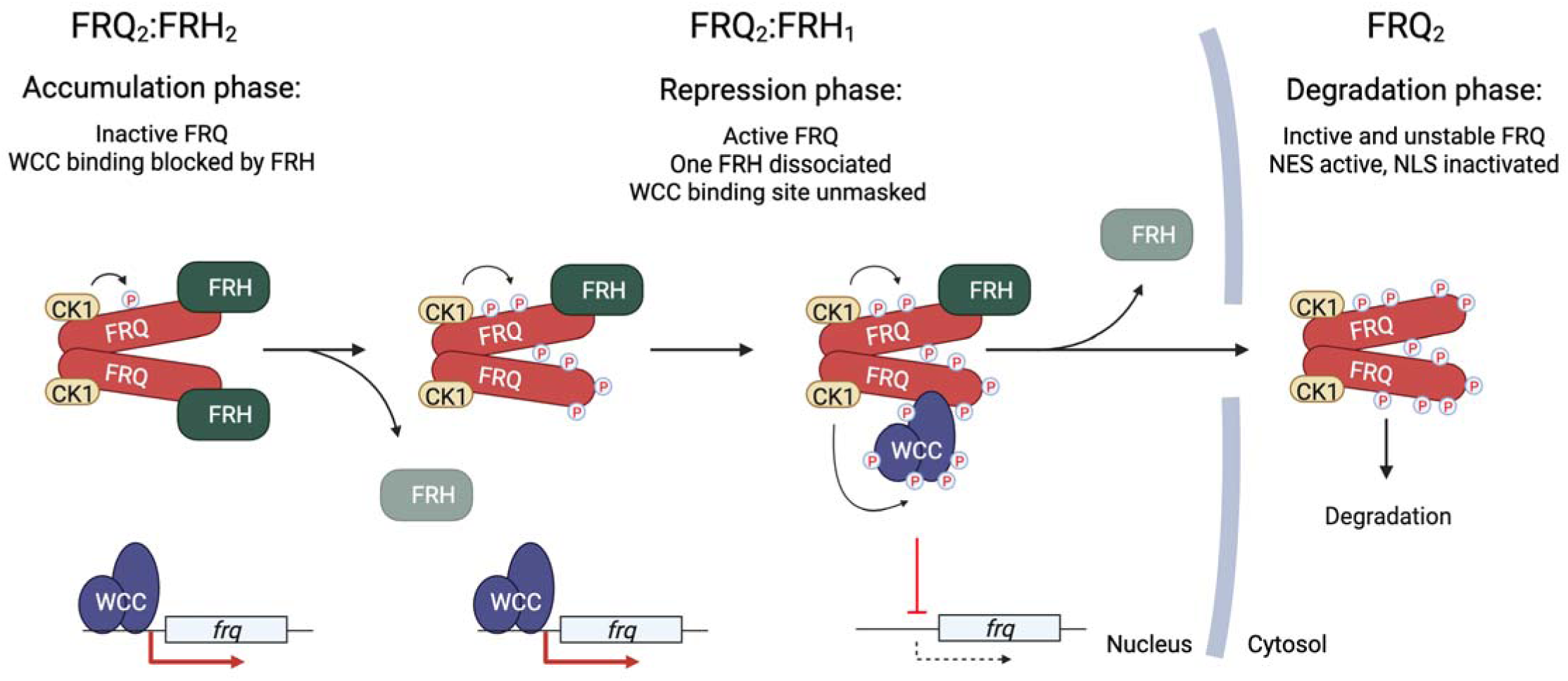
Model of CK1a-dependent subunit remodeling in the FRQ-FRH complex. Newly synthesized, unphosphorylated FRQ accumulates with FRH in the nucleus as a heterotetramer with an FRQ_2_FRH_2_ stoichiometry. The nuclear accumulation of FRQ is facilitated by its three nuclear localization signals (NLSs) and is further supported by a tight association with FRH, which provides a nuclear import signal and may additionally mask FRQ’s nuclear export signal (NES). The two bound FRH molecules block the WCC binding sites on FRQ, rendering the FRQ_2_FRH_2_ complex inactive. Consequently, the WCC remains active, promoting *frq* transcription and supporting the accumulation of high levels of inactive FRQ_2_FRH_2_. Phosphorylation of FRQ inactivates its NLSs and, after a considerable delay, triggers the dissociation of FRH. Due to the slow and partially stochastic nature of FRQ phosphorylation, the release of the two FRH molecules from a FRQ dimer is not synchronous, leading to the transient formation of a heterotrimeric FRQ_2_FRH_1_ species. The FRQ_2_FRH_1_ complex is the active species because it exposes a WCC binding site, allowing CK1a to phosphorylate and inactivate transiently interacting WCCs, thereby shutting down *frq* transcription. The FRQ_2_FRH_1_ complex can be exported via the exposed NES in one FRQ subunit but will be reimported via the FRH bound to the second FRQ subunit. Further phosphorylation of FRQ by CK1a eventually triggers the dissociation of the second FRH molecule, resulting in an FRQ_2_-only species. At this stage, FRQ’s NLSs are fully inactivated by phosphorylation, prompting relocalization of FRQ_2_ to the cytosol, where it can no longer support phosphorylation of nuclear WCC and is eventually degraded, thus closing the negative feedback loop. Subsequent dephosphorylation of WCC initiates a new circadian cycle. Created with <u>BioRender.com</u>.

Our model not only aligns with previous findings but also offers new mechanistic insights into previously unresolved mutant phenotypes. For instance, we found that the FRH^R806H^ mutant protein blocks the WC-1 binding site more effectively than wild-type FRH. This increased binding affinity likely accounts for the absence of WCC-FRQ interaction reported in *Neurospora* strains expressing FRH^R806H^ ^28^. Additionally, monomeric mutants of FRQ, while able to interact with FRH and CK1, remain nonfunctional ^3,25,26^. According to our model, monomeric FRQ in complex with FRH stays inactive due to its inability to bind WCC. After the phosphorylation-dependent release of FRH, monomeric FRQ exposes its WCC binding site and should be active in negative feedback. However, in the absence of a bound FRH, it will be rapidly exported and degraded in the cytosol, similar to FRQ^6B2^, preventing it from accumulating in sufficient quantities to exert negative feedback in the nucleus. The mutations in the N-terminal region of FRQ, previously reported to impair WCC binding ^27^, fall within the predicted dimerization domain and therefore are more likely to affect FRQ dimerization than its direct interaction with WCC.

Several aspects of our model require further validation and refinement in future work. One question is whether the physiologically relevant form of FRQ in *Neurospora* is a dimer or a higher-order oligomeric assembly, and how these forms may be regulated. The nuclear-cytoplasmic distribution of FRQ appears to be influenced by the number and strength of its NLSs relative to NES. While we have demonstrated that an FRQ monomer contains three NLSs and one NES regulated by phosphorylation, further investigation is required to elucidate the precise mechanism by which phosphorylation inactivates the NLSs. Because the polybasic NLSs are located near multiple phosphorylation sites, their gradual inactivation may result from charge neutralization. It is also essential to explore how phosphorylation of FRQ weakens the interaction with FRH to expose FRQ’s NES and the WCC binding site. Perhaps the most compelling and exciting direction for future research, facilitated by the cell-based system established here, is to investigate how FRQ and CK1a influence the subcellular dynamics and chromatin association of the WCC.

While our model will undoubtedly evolve with future discoveries, it provides a detailed conceptual framework for understanding how this eukaryotic circadian clock measures time at the molecular level and offers an additional experimental system for further research in this field.

## Methods

### Plasmids

The N-terminal mKATE-2 (mK2) tag, derived from pmKate2-N and the C-terminal mNeonGreen (mNG) tag, derived from pMaCTag-07 ^30^, were inserted into pcDNA™4/TO downstream of an inhibitory TetR-responsive cytomegalovirus (CMV-TetO) promoter by cloning with overlapping regions using appropriate primers ^31^. In short, a linear vector and an insert DNA fragment carrying a 30 bp sequence overlap were transfected and recombined *in vivo* in *E. coli*. Accordingly, the cDNAs of the *Neurospora crassa* genes, *frh*, *ck1a*, *frq* and fragments of *frq* were inserted into pcDNA™4/TO, pcDNA™4/TO-mK2 and pcDNA™4/TO-mNG as indicated. The 6B2 mutation as well as the premature stop codon of the FRQ^9^ mutant were introduced in the pcDNA™4/TO-mK2-*frq* by site directed mutagenesis using the QuikChange II kit (Agilent).

FRQ-NLS mutants for expression in *Neurospora* were introduced by site directed mutagenesis into pBM60-*frq*, resulting in pBM60-*frq^mNLS1^*, *frq^mNLS2^,* -*frq^mNLS3^* and -*frq^mNLS1/2/3^* (QuikChange II kit, Agilent).

### Western blots

Western blotting was performed as previously described ^32^.Chemiluminescence signals were detected using X-ray films, which were then developed in Konica Minolta SRX-101A Medical Film Processor.

### U2OS T-REx cells and culture conditions

U2OS T-REx (U2OStx) cells were maintained in a 5% CO_2_ incubator at 37°C in Dulbecco’s modified Eagle’s medium (DMEM, Thermo Fisher Scientific) supplemented with 10% fetal bovine serum (FBS, Thermo Fisher Scientific), 1% penicillin-streptomycin (Thermo Fisher Scientific) and Hygromycin B (50 μg/mL, Invivogen).

Leptomycin B (Merck; CAT# L2913-.5UG) was added at a final concentration of 20 nM one hour after doxycycline (DOX) induction.

### Protein expression

U2OStx cells were seeded in 96 well plates 24 hours prior to transfection (Corning, CAT#3598). A fixed amount of plasmid DNA (150 or 200 ng as indicated), pcDNA™4/TO plasmids with the genes of interest balanced with empty vector, was transfected per well using Xfect™ Transfection Reagent (Takara Bio) according to the manufacturer’s instructions. Protein expression was induced 24 hours post transfection by addition of 10 ng/mL DOX. pcDNA4/TO-FRQ-mNG was co-transfected pcDNA4/TO-FRQ at a ratio of 25 ng : 50 ng to avoid overexposure of mNG fluorescence.

### Protein extraction for Western blot

Cells were seeded in 12-well plates 24 hours before transfection. At either 4 or 14 hours after DOX induction, cells were harvested directly in 2x protein sample buffer and boiled at 95°C for 7 minutes before Western blot analysis.

### Live-cell microscopy

After induction with DOX, 96-well plates were placed in either the Incucyte ZOOM® or Incucyte® SX1 system (Sartorius) at 37°C. Images were acquired at 1-hour intervals for 48 hours and then analyzed using the Incucyte® 2023A GUI (Sartorius) and Incucyte® 2016B GUI (Sartorius) software.

### Quantification and statistical analysis

IncuCyte analysis: In each experiment, 20 to 30 cells were quantified. Cells with fluorescence levels that were either too low or excessively high were excluded from the analysis. The cytoplasmic and nuclear distributions of mNG- and mK2-tagged proteins were examined, along with their colocalization and the number of foci as specified in each respective experiment. WC-1-mNG was considered as localizing to the nucleus when nuclear green fluorescence intensity was higher than in the cytosol. Interaction between FRQ and WC-1 was defined by either co-localization within the same nuclear foci or homogeneous nuclear co-localization, as opposed to segregation into separate foci or cytosolic WC-1 alongside nuclear FRQ. Data were visualized using BioRender Graph (Created in https://BioRender.com).

### *Neurospora crassa* culture conditions

Conidial suspensions in 1 M sorbitol were prepared from strains grown (5–7 days) on standard solid growth medium (2.2% agar, 0.3% glucose, 0.17% L-arginine, 1x Vogel’s medium and 0.1% biotin). Standard growth medium for liquid cultures contained 2% glucose, 0.17% L-arginine and 1x Vogel’s medium. Liquid cultures were inoculated with conidia and grown in constant light at 25 °C for 2 to 3 days if not indicated otherwise.

To express hypophosphorylated FRQ, 48 h light-grown liquid cultures were transferred to the dark for 10 hours (essentially leading to the degradation of “old” FRQ) and then exposed to light for 3 hours to induce a burst of FRQ expression before mycelia were harvested and native protein extract was prepared.

### *Neurospora* strains

**Table 1.**
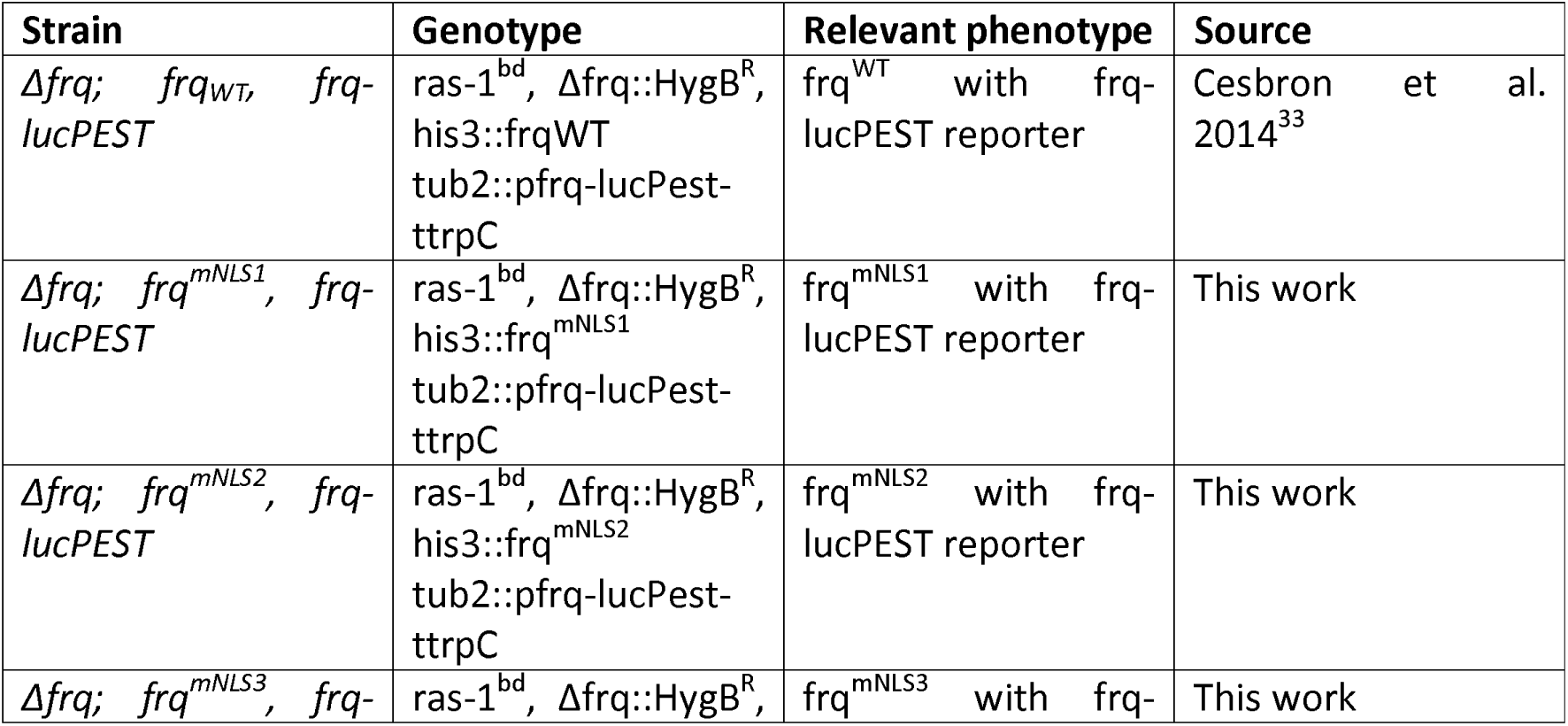
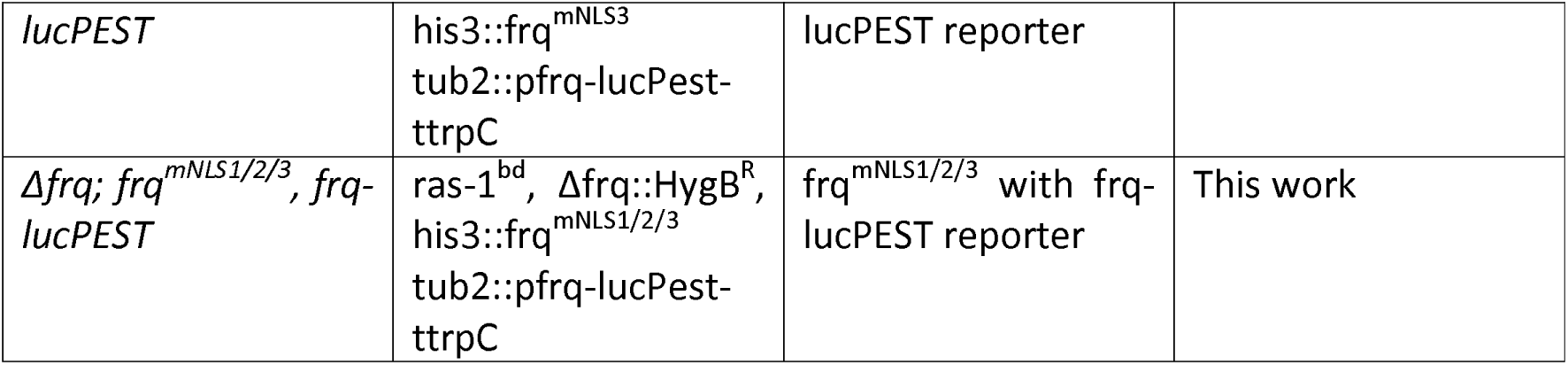
*Neurospora crassa* strains used in this study.

| Strain | Genotype | Relevant phenotype | Source |
| --- | --- | --- | --- |
| $\Delta frq$; $frq_{WT}$, $frq$-<br>*lucPEST* | $ras-1^{bd}$, $\Delta frq::HygB^R$,<br>$his3::frq_{WT}$<br>$tub2::pfrq$-*lucPest*-<br>*ttrpC* | $frq^{WT}$ with $frq$-<br>*lucPEST* reporter | Cesbron et al.<br>2014<sup>33</sup> |
| $\Delta frq$; $frq^{mNLS1}$, $frq$-<br>*lucPEST* | $ras-1^{bd}$, $\Delta frq::HygB^R$,<br>$his3::frq^{mNLS1}$<br>$tub2::pfrq$-*lucPest*-<br>*ttrpC* | $frq^{mNLS1}$ with $frq$-<br>*lucPEST* reporter | This work |
| $\Delta frq$; $frq^{mNLS2}$, $frq$-<br>*lucPEST* | $ras-1^{bd}$, $\Delta frq::HygB^R$,<br>$his3::frq^{mNLS2}$<br>$tub2::pfrq$-*lucPest*-<br>*ttrpC* | $frq^{mNLS2}$ with $frq$-<br>*lucPEST* reporter | This work |
| $\Delta frq$; $frq^{mNLS3}$, $frq$- | $ras-1^{bd}$, $\Delta frq::HygB^R$, | $frq^{mNLS3}$ with $frq$- | This work |
| *lucPEST* | his3::frq<sup>mNLS3</sup><br>tub2::pfrq-lucPest-<br>ttrpC | lucPEST reporter |  |
| Δfrq; frq<sup>mNLS1/2/3</sup>, frq-<br>*lucPEST* | ras-1<sup>bd</sup>, Δfrq::HygB<sup>R</sup>,<br>his3::frq<sup>mNLS1/2/3</sup><br>tub2::pfrq-lucPest-<br>ttrpC | frq<sup>mNLS1/2/3</sup> with frq-<br>lucPEST reporter | This work |

### Native protein extraction

To extract native protein from *Neurospora*, mycelial tissue was ground in liquid nitrogen using a precooled mortar and pestle. The resulting powder was suspended in an extraction buffer containing 50 mM Hepes-KOH (pH 7.4), 137 mM NaCl, 10% (v/v) glycerol, and 5 mM EDTA, with 1 mM phenylmethylsulfonyl fluoride (PMSF), leupeptin (5 μg/ml), and pepstatin A (5 μg/ml). Protein concentration was measured using a NanoDrop (PeqLab).

### Luciferase reporter assay

The luciferase assay was performed as described previously ^33^ with slight modifications: 96-well plates containing growth medium including 75 µM D-lucferin were inoculated with 3×10^4^ conidia per well and incubated in constant darkness at 25°C for 3 days. The plate was then transferred to an incubator at 25 °C with an EnSpire Multilabel Reader (Perkin Elmer). Bioluminescence was measured in 30 min intervals. The following light/dark regime was used: 2 h darkness, 12 h light, 12 h darkness, 12 h light, constant darkness. Light intensity used was 30 µE. Data was normalized to average bioluminescence levels during light exposure.

### *In vitro* phosphorylation

For *in vitro* phosphorylation of FRQ, *Neurospora* whole cell lysate was incubated over night at 4 °C with 2 µg purified recombinant CK1a ^16^ per mg of lysate in a buffer containing a final concentration of 50 mM Hepes/KOH pH 7.4, 120 mM NaCl, 11.3 mM MgCl_2_, 12.5 mM ATP, 1xPhosStop (Roche), 1 mM PMSF, 1 µg/ml leupeptin, 1 µg/ml pepstatin.

### Co-immunoprecipitation

Co-IP was performed as described ^26^. Briefly, 20 µl of M2 FLAG Sepharose beads (Sigma) were washed twice in PBS. 4 mg of whole cell lysate from a strain expressing N-terminally 2xFLAG-tagged FRH was then added to the beads and filled to 500µl with PBS supplied with protease inhibitors (Roche) and incubated for 3 h at 4 °C. Supernatant was removed, beads washed twice and bound protein eluted by boiling at 95 °C for 5 minutes in 2x Laemmli. 400 µg of the input and supernatant were loaded together with amounts of IP corresponding 1x (for FLAG/FRH decoration) and 9x (for FRQ decoration) equivalents of the input. Western blots were probed with antibodies against FRQ and FLAG (FRH).

### Size exclusion chromatography

A Superose 6 Increase G10/300 GL column was used in an Äkta Pure system (GE Life Sciences/Cytiva). The column was equilibrated with gelfiltration buffer (25 mM HEPES/ KOH (pH 7.4), 140 mM NaCl, 1 mM EDTA pH 8.0, 1% glycerol, 0.05% Triton X-100). 10 mg native *Neurospora* protein extracts were loaded onto a 200 µL loop and chromatography was performed at 4°C with a flow rate of 0.5 mL/min. Fractions of 490 µL were collected, and protein was precipitated by adding 122.5 µL of 50% TCA (w/v). The samples were incubated on ice for 10 minutes and then centrifuged (4°C, 14,000 g, 10 minutes). Pellets were washed with 1 mL acetone, air-dried for 3 minutes at room temperature, and resuspended in 90 µL of 2x Laemmli buffer. Samples were boiled at 95°C for 5 minutes. 45 µL was loaded on SDS-PAGE, followed by Western blotting for analysis.

## Author contribution

CS: U2OStx Incucyte experiments

BR: some U2OStx Incucyte experiments

SS & ACRD: *Neurospora* experiments

MB: Conceptualization, writing.

## Acknowledgements

We thank Alessia Ruggieri und Michael Knop for providing mKATE-2 and mNeonGreen templates, respectively. The work was supported by the Deutsche Forschungsgemeinschaft, TRR186.

## Supplemental Fig. legends

**Fig. S1:**
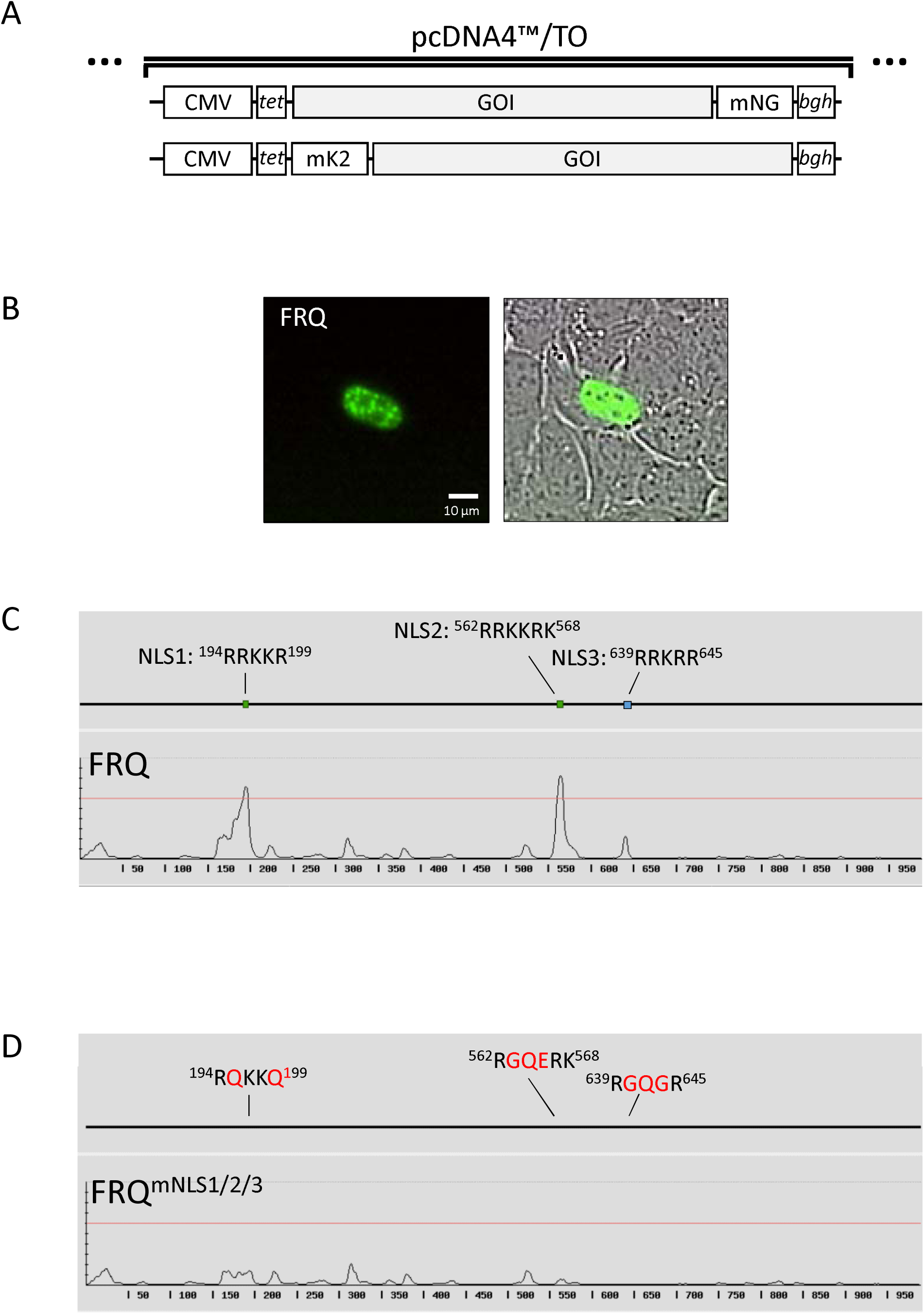
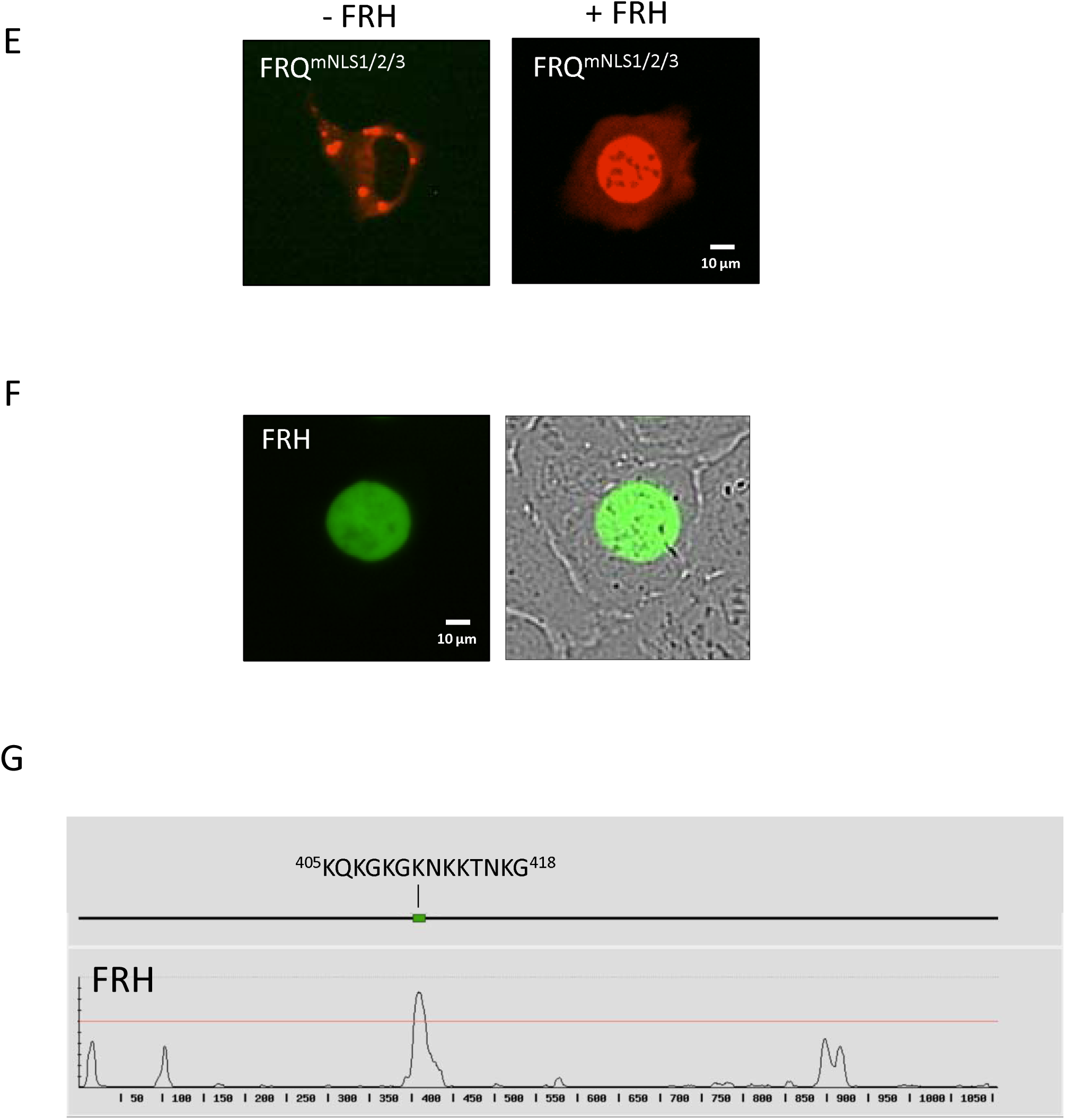
**A** Schematic of pcDNA expression vectors. The gene of interest (GOI) is expressed under the control the CMV promoter followed by a TET operator. It is tagged either with a C-terminal mNeonGreen (mNG) or an N-terminal mKate2 (mK2) followed by the 3‘-UTR of the *Bovine Growth Hormone* Gene, *bgh*. **B** Transient expression of FRQ-mNG in U2OStx cells. **C** Prediction of NLSs in FRQ (http://www.moseslab.csb.utoronto.ca/NLStradamus/). **D** Prediction of NLSs in FRQ^mNLS1/2/3^. Mutations introduced in NLS1, NLS2 and NLS3 of FRQ are indicated by red letters. **E** Subcellular localization of mK2-FRQ^mNLS1/2/3^ left panel: without FRH, right panel: with FRH-mNG (not shown). **F** FRH-mNG is localized in the nucleus of U2OStx cells. **G** Prediction of NLSs in FRH. The predicted NLS (aa 405-418) is not resolved in the crystal structure (PDB:4XGT) and predicted to be unfolded (AF: Q873J5-F1).

**Fig. S2:**
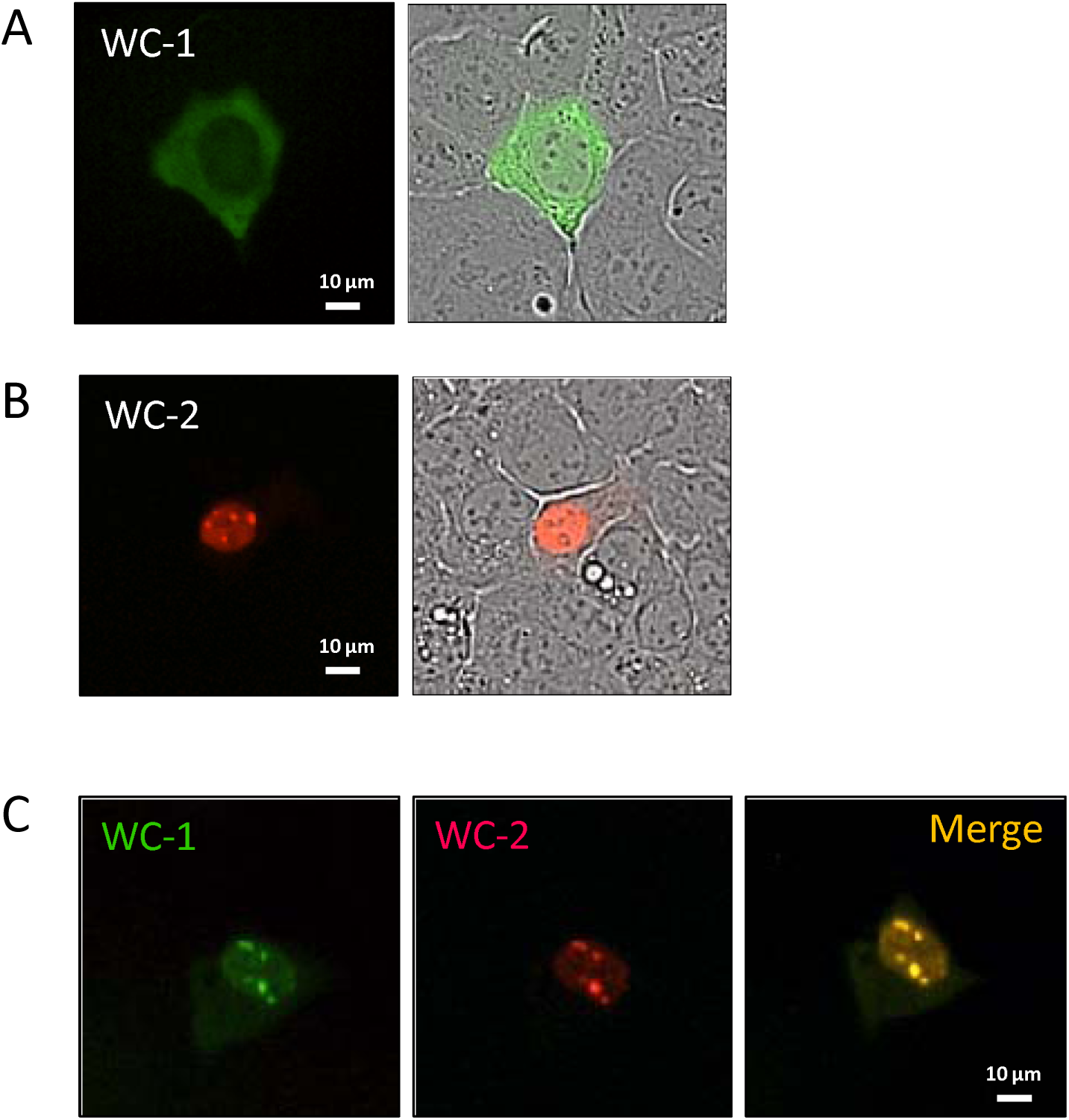
Expression and subcellular localization of WC-1 and WC-2 in U2OStx cells. **A** WC-1-mNG accumulates in the cytosol. **B** mK2-WC-2 forms nuclear foci. **C** Upon co-expression, WC-1-mNG and mK2-WC-2 accumulate in nuclear foci.

**Fig. S3:**
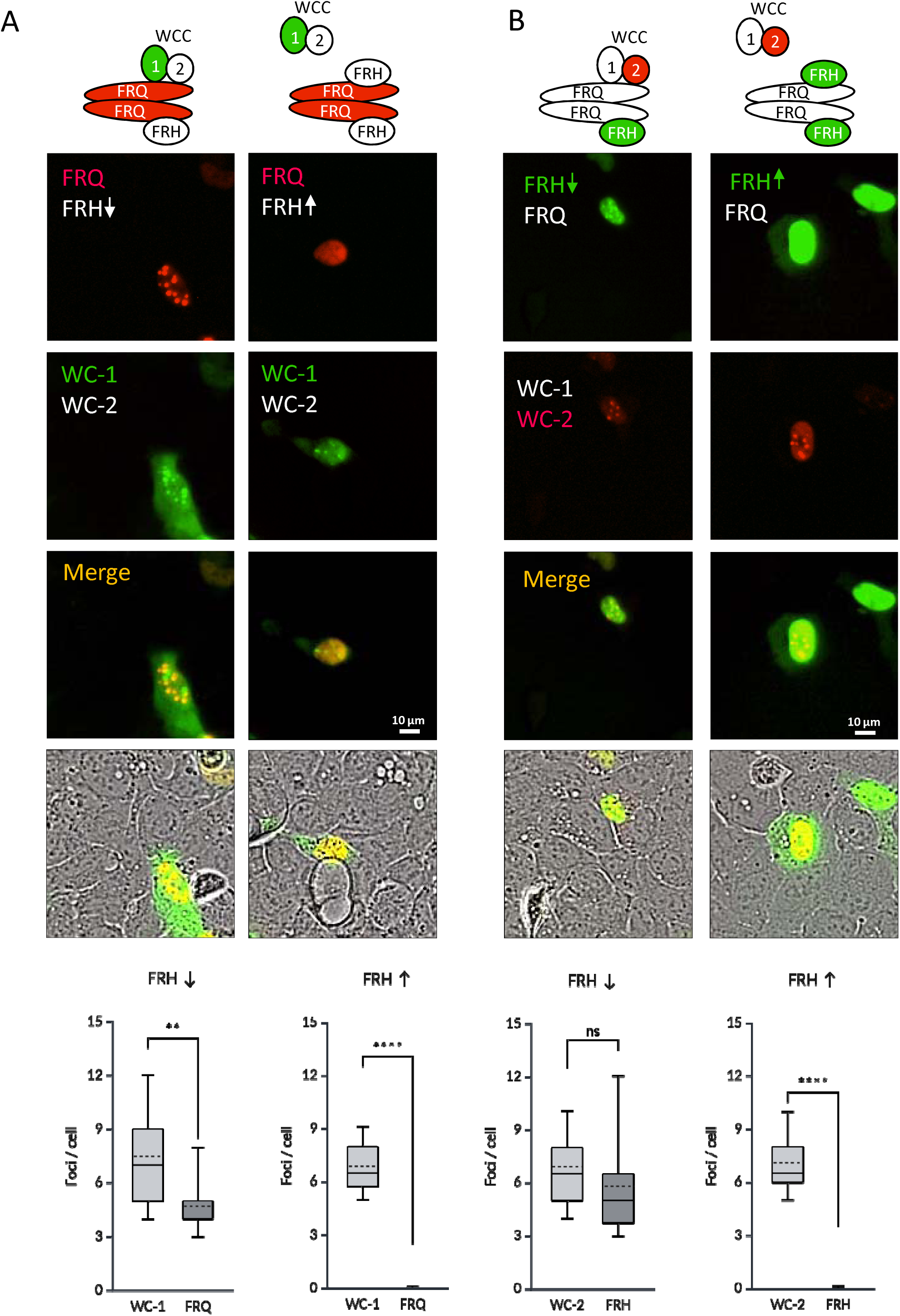
Saturating amounts of FRH interfere with binding of FRQ to WCC. Top: Schematics depict the interactions of fluorescently tagged (green and red) and untagged (white) proteins. **A** Left: Co-expression of mK2-FRQ (50 ng) and sub-saturating amounts of untagged FRH (12.5 ng) with WC-1-mNG (50 ng) and untagged WC-2 (25 ng). mK2-FRQ and WC-1-mNG co-localize in nuclear foci. Right: Co-expression of mK2-FRQ (40 ng) and saturating amounts of untagged FRH (80 ng) with WC-1-mNG (50 ng) and untagged WC-2 (25 ng). mK2-FRQ is dispersed in the nucleus. **B** Left: Co-expression of untagged FRQ (50 ng) and sub-saturating amounts of FRH-mNG (12.5 ng) with untagged WC-1 (50 ng) and mK2-WC-2 (25 ng). FRH-mNG and mK2-WC-2 co-localize in nuclear foci. Right: Co-expression of mK2-FRQ (40 ng) and saturating amounts of untagged FRH (80 ng) with WC-1-mNG (50 ng) and untagged WC-2 (25 ng). FRH-mNG is dispersed in the nucleus and mK2-WC-2 -mNG accumulates in nuclear foci. Bottom: Quantification of nuclear foci of WC-1-mNG and mK-FRQ A and FRH-mNG and mK-WC2 B, respectively. Unpaired t-test, **: p ≤0.01, ****: p ≤0.0001, ns: not significant p =0.0691.

**Fig. S4:**
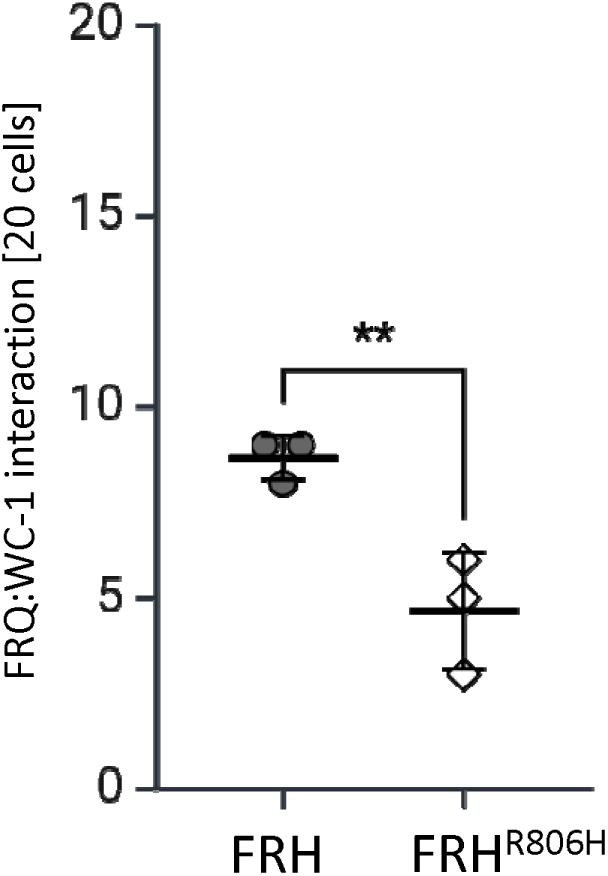
FRH^R806H^ competes better than FRH with the binding of WC-1 to FRQ. Co-expression of mK2-FRQ (40 ng) and WC-1-mNG (50 ng) with non-saturating amounts (20 ng) of either untagged FRH or FRH^R806H^. Interaction of mK2-FRQ with WC-1-mNG was evaluated by counting cells displaying nuclear enrichment of WC-1-mNG. 20 cells each from three independent experiments were evaluated. Unpaired T-test, **: p ≤ 0.01.

**Fig. S5:**
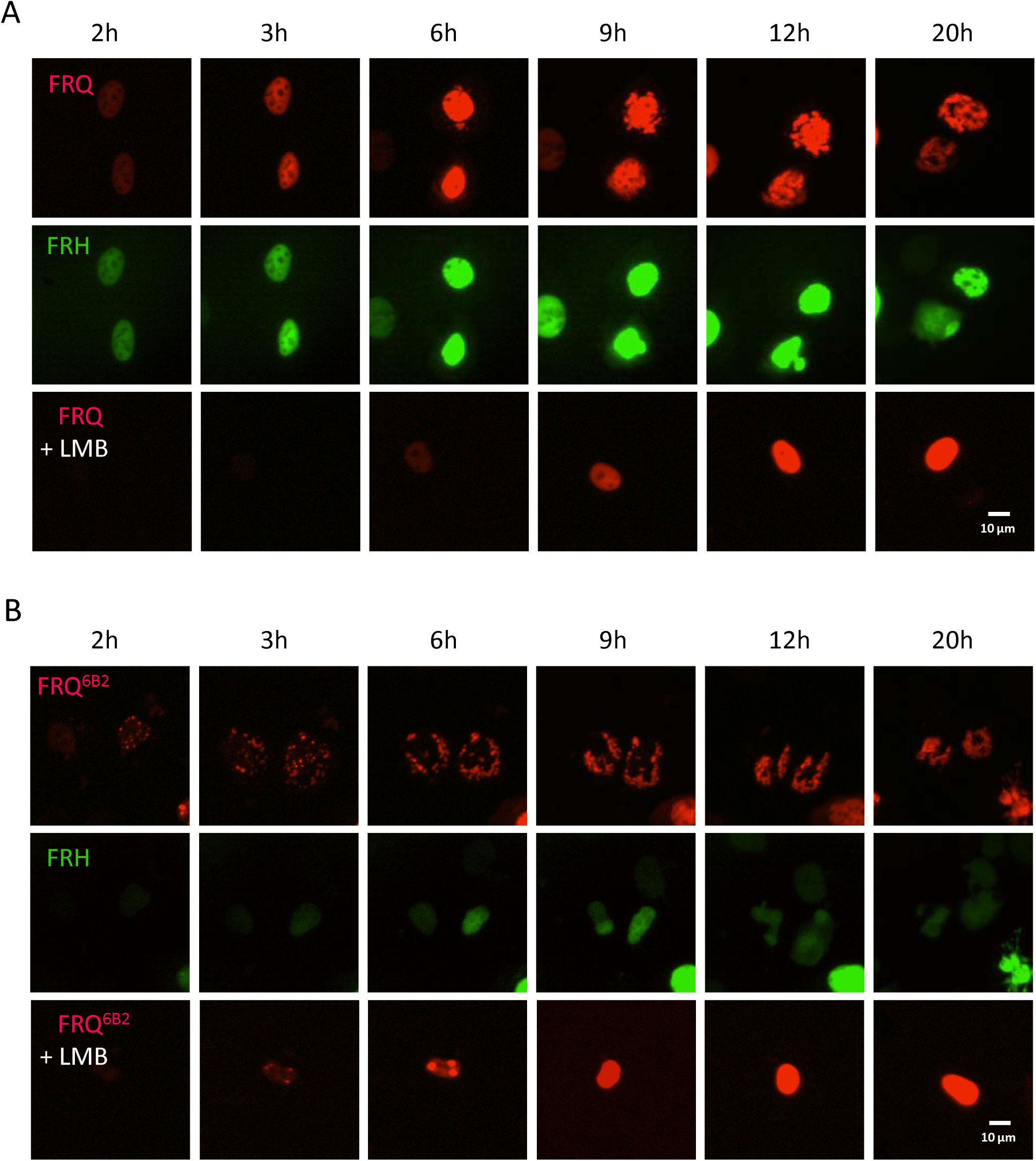
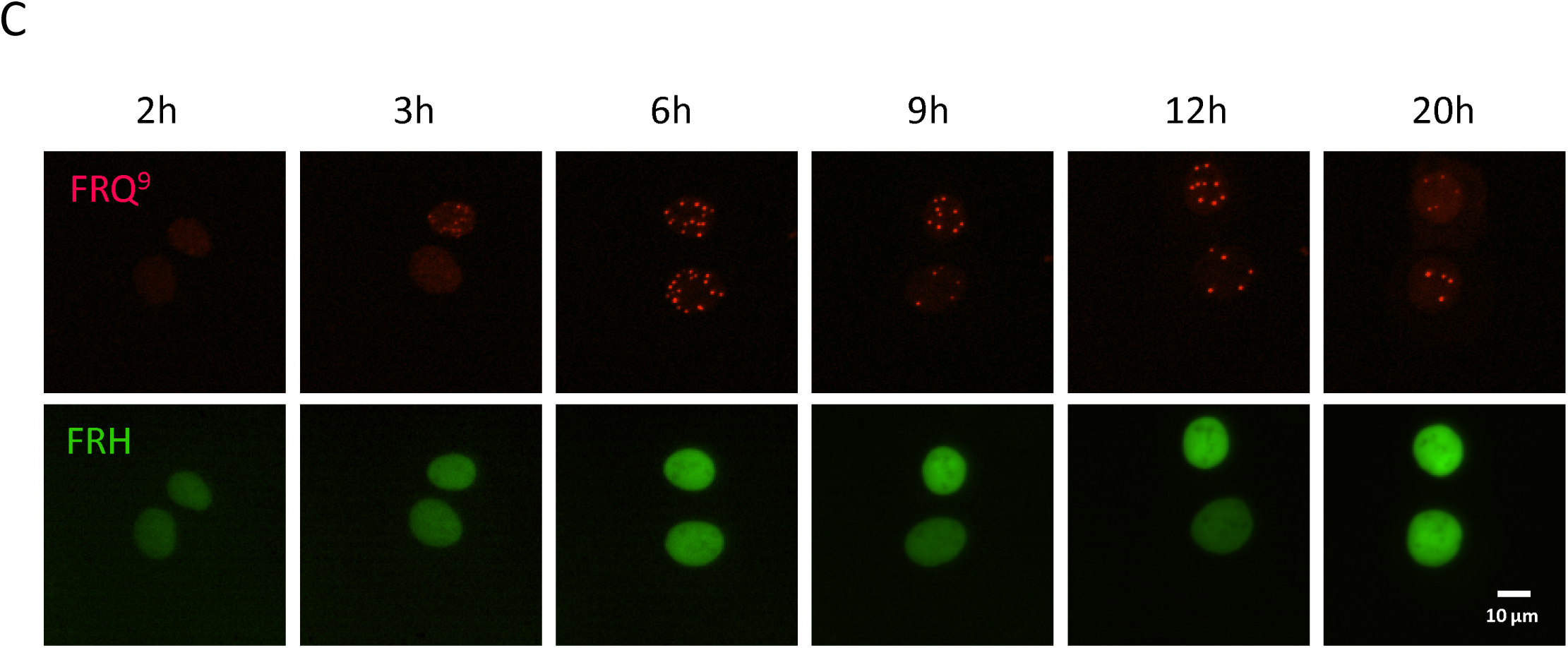
Phosphorylation of FRQ by CK1a triggers its dissociation from FRH and nuclear export. **A - C** Co-expression of **A** mK2-FRQ, **B** mK2-FRQ^6B2^ or **C** mK2-FRQ^9^ with FRH-mNG and untagged CK1a for the indicated time periods. LMB was added when indicated. Data are identical to Fig. 6 but show separate panels for the mK2-tagged FRQ versions and FRH-mNG.

**Fig. S6:**
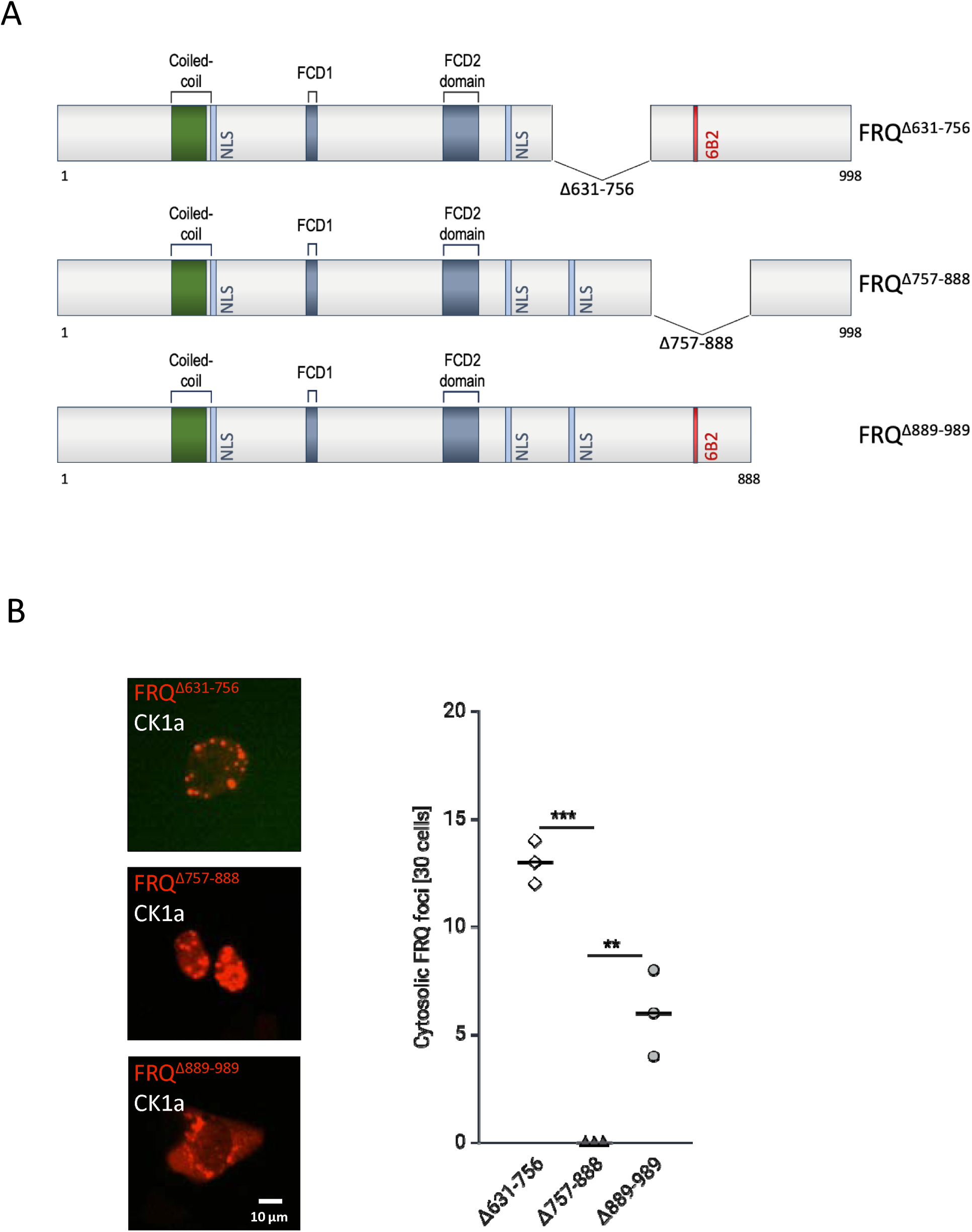
FRH promotes nuclear import of FRQ and masks FRQ’s NES. **A** Schematic of FRQ deletion constructs. The proteins were tagged N-terminally with mK2. **B** Subcellular localization of mK2-FRQ^Δ631–756^ mK2-FRQ^Δ757-888^ and mK2-FRQ^Δ889-989^ co-expressed with untagged CK1a in U2OStx cells. Graph shows quantification of cells with cytoplasmic localization of mutant mK2-FRQ. 30 cells each from three independent experiments were evaluated. Unpaired T-test, **: p ≤ 0.01, ***: p ≤ 0.001.

**Fig. S7:**
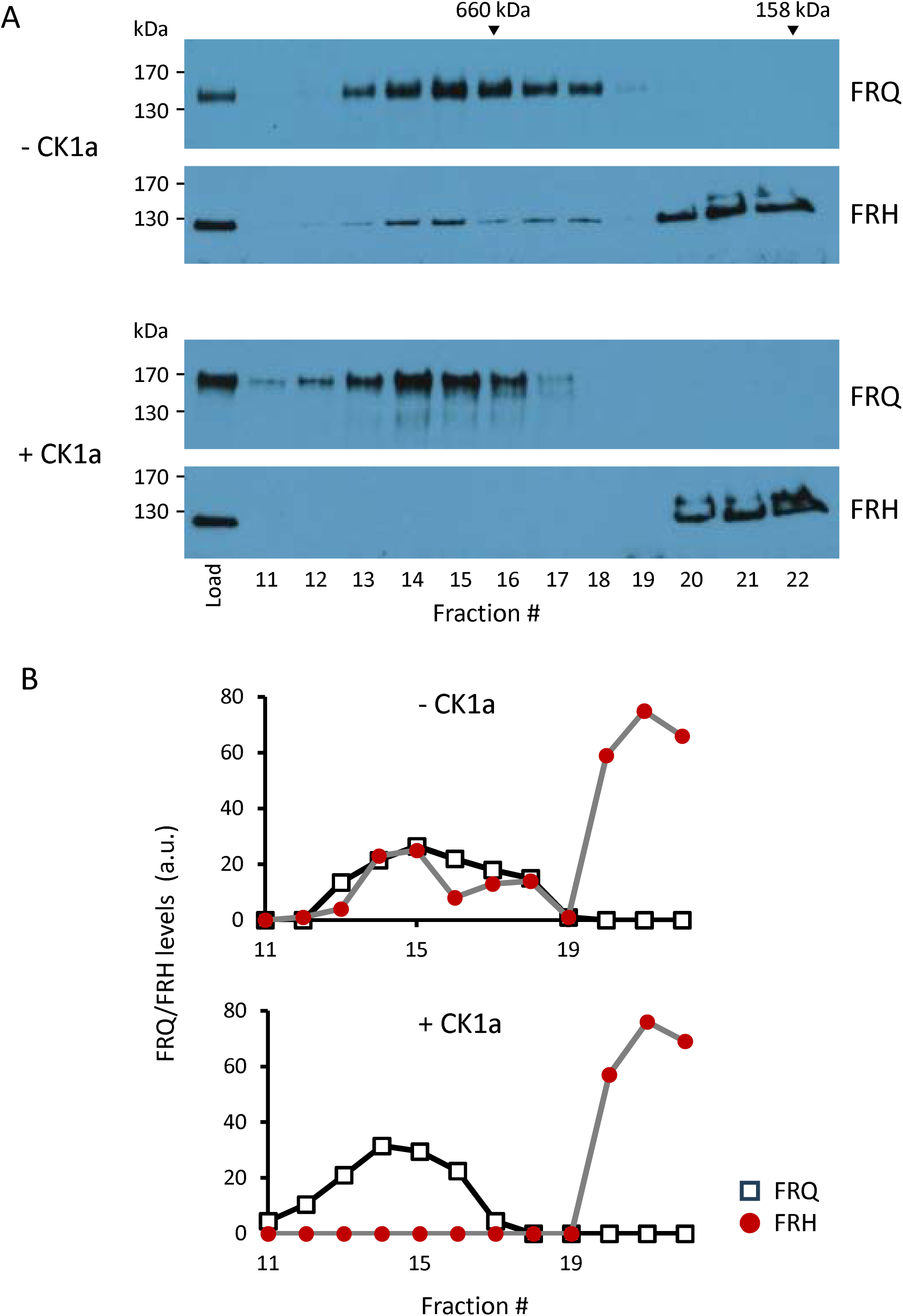
Size exclusion chromatography of FRH and hypo- and hyper-phosphorylated FRQ, respectively. **A** Whole cell lysate from *Neurospora* expressing hypophosphorylated FRQ was treated with or without CK1a. The samples were then analyzed by size exclusion chromatography using a Superose 6 Increase 10/300 GL column. FRQ and FRH were detected by Western blot. **B** Western blots were quantified using ImageJ. Densitometric signals were plotted against the fraction number.

